# Myeloid cells contribute to bystander CD8 T cell accumulation in metabolic-associated steatohepatitis and are sufficient for fibrosis

**DOI:** 10.64898/2025.12.17.694415

**Authors:** Cynthia Lebeaupin, Katelyn L. Donahue, Samir Lal, Stephen M. Christensen, Shoh Asano, Marc H. Wadsworth, Kathryn Bound, Saurav De, James McMahon, Franklin Schlerman, Chang Wang, Xiao Chen, Alexander M.S. Barron, Thomas A. Wynn, Kevin M. Hart, Thomas Fabre

**Author notes:** Corresponding author 1 Portland Street, Cambridge MA, USA. These authors contributed equally to this work as first authors (Cynthia Lebeaupin, Katelyn L. Donahue).

## Abstract

**Background & Aims:** Metabolic dysfunction-associated steatohepatitis (MASH) is a leading cause of liver fibrosis, morbidity, and mortality. It is characterized by the accumulation of fat in hepatocytes causing cell death followed by stromal cell activation and scar deposition at later disease stages. While a marked accumulation of cytotoxic CD8 T cells is observed in MASH in humans and mice, the role of adaptive immune cells in fibrosis progression remains debated.

**Methods:** Transcriptional data were curated from human datasets and preclinical mouse models, including mice deficient in TCRαβ T cells (*Tcrb^-/-^*), which lack conventional CD4 helper and CD8 cytotoxic T cells, in diet-induced fibrosing MASH for single-cell RNA-sequencing. Specific human and mouse liver transcriptomic immune signatures were corroborated by flow cytometry and immunofluorescence.

**Resultsx:** MASH was associated with the expansion of lymphocytes and myeloid cells. Auto-aggressive CXCR6^+^PD1^+^FASLG^+^ CD8 T cells were enriched in livers of high-fat-diet-fed mice and correlated with MASH and fibrosis. Notably, *Tcrb-*deficient mice in a chemical and dietary preclinical MASH model developed fibrosis to the same extent as wild-type mice and exhibited exacerbated pro-fibrotic type 3 inflammation. Loss of conventional CD8 T cells neither impacted myeloid cell number nor phenotype within the fibrotic niche. Neutrophil- and non-conventional lymphocyte-derived GM-CSF and IL-17A were central drivers in the fibrotic niche composed of pathogenic macrophages and activated myofibroblasts. Furthermore, myeloid cells were identified as the main source of CXCL16 in diseased livers, retaining auto-aggressive CD8 T cells to the scar and contributing to their accumulation. Thus, non-adaptive cells, including scar-associated macrophages (SAMs) and myofibroblasts, are sufficient for MASH-driven fibrosis in this model.

**Conclusions:** Targeting myeloid cells and fibroblasts should be prioritized as anti-fibrotic therapies for MASH.

**Graphical abstract:** The immunopathological niche in fibrosis

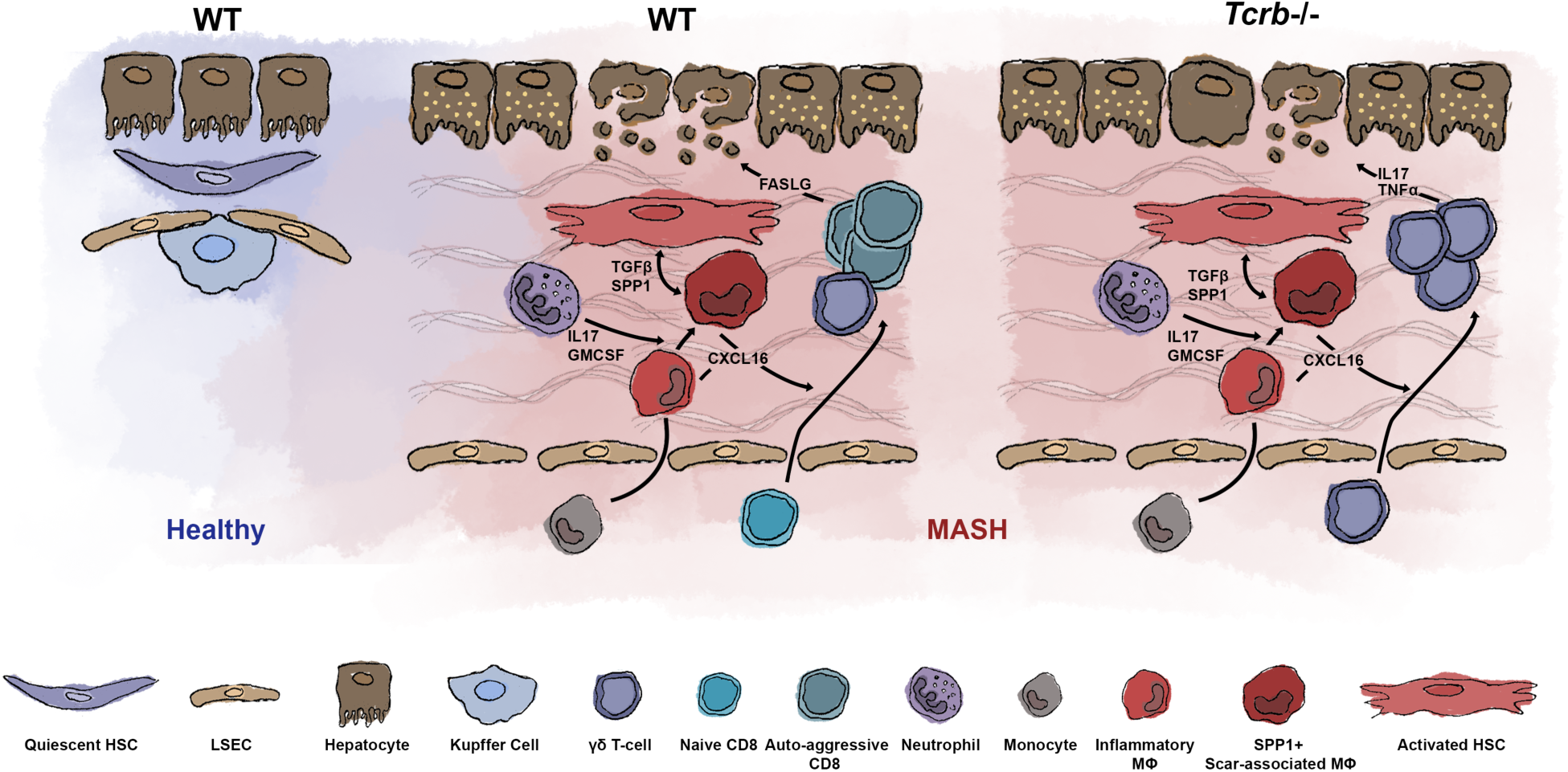

**Highlights:** Adaptive and innate immune cells localize to the human MASH fibrotic niche

CD8 T cells increase with MASH severity while MHC-II^+^ cells accumulate with fibrosis

CXCR6^+^FASLG^+^ CD8 auto-aggressive T cells accumulate at the scar

CXCL16^+^ scar-associated macrophages retain CD8 T cells in the fibrotic MASH niche

Liver injury, inflammation and fibrosis persist in the absence of conventional T cells

**Impact and implications:** Metabolic dysfunction-associated steatohepatitis (MASH) is a progressive liver disease in which fibrosis is a key determinant of patient outcomes and survival. This study demonstrates that fibrosis in MASH can develop independent of conventional T cell immunity, driven by myeloid cells including scar-associated macrophages (SAMs) and neutrophils within a localized hepatic niche in mice and humans. This highlights the need to prioritize innate immune mechanisms as key drivers of chronic (late-stage) liver disease for researchers and clinicians developing anti-fibrotic therapies.

## Introduction

Fibrosis in metabolic dysfunction-associated steatohepatitis (MASH) represents a critical and unresolved clinical challenge of chronic liver disease that will continue to rise over the next three decades^1^. While recent advances in metabolic therapies show promise in reducing steatosis and hepatocellular injury, fibrosis often persists^2,3^. MASH with fibrosis perpetuates due to a conserved, self-perpetuating cycle of injury, inflammation, cell differentiation, and scarring^4^. A critical factor in fibrotic progression and maintenance is the initiating inflammatory cascade. This involves the recruitment and differentiation of innate immune cells (*e.g.,* monocytes to macrophages, neutrophils) as well as the recruitment and activation of adaptive immune cells (*e.g.,* T cells), leading to the establishment of a complex fibrotic niche^4^.

Historically, the role of adaptive immunity in fibrosis has been contentious due to discrepancies between preclinical models and human diseases. The involvement of adaptive immune cells (*e.g.,* T and B cells) in viral hepatitis (HBV, HCV) and auto-immune hepatitis in humans is evident^5–10^. Auto- or viral-antigen-specific adaptive cells contribute to hepatocellular damage and activation of myofibroblasts via both cytokine secretion and canonical cell death mechanisms (*e.g.,* perforin/granzyme)^11–14^. In contrast, preclinical studies using carbon tetrachloride (CCl_4_) or dimethylnitrosamine (DMN)-induced liver injury in SCID and NOD/SCID mice demonstrated that adaptive immunity was dispensable for fibrosis development^15,16^. However, the hepatotoxicity and non-physiological mode of acute “sterile” chemical preclinical injury, such as CCl_4_ and DMN, limits the translational relevance and support of these findings. Moreover, the absence of adaptive immunity in these settings may have altered the kinetics and quality of innate immune responses. Despite these limitations, these foundational studies laid the groundwork for a broader understanding of fibrosis as a maladaptive wound-healing response, potentially uncoupled from classical adaptive immune pathways^17^. In the context of MASLD/MASH, perceived originally as a “sterile” injury, more recent studies reported the expansion of Th2, Th17, Tregs and CD8 T cells in liver biopsies, potentially resulting from dysbiosis and inflammation related to the peripheral metabolic syndrome^11,18,19^.

Investigations of adaptive immunity using diet-induced preclinical MASH models benefit from a more organic, less “sterile” injury setting. In methionine and choline-deficient diet (MCDD)-induced MASH, the ablation of XCR1^+^ dendritic cells (DCs) decreased T cell activation and reduced liver injury in mice^20^. In another report, choline-deficient high-fat diet (CDHFD) induced the metabolic activation of CD8 T cells and NKT cells that promoted liver damage and MASH^19^. Recently, auto-aggressive CXCR6⁺PD1⁺FASLG⁺ CD8 T cells that activate in response to calcium influx rather than antigen recognition were shown to accumulate in high-fat diet- (HFD)-driven MASH as well as human fibrotic livers. These auto-aggressive CD8 T cells exhibit transcriptional profiles reminiscent of pathogenic T cells in chronic inflammatory diseases and mediate FAS/FASLG-induced epithelial cell death^21^. However, these HFD models only showed modest fibrosis induction and limited human validation. Furthermore, the contribution of auto-aggressive T cells to fibrosis remains correlative, as the focus was on the composite non-alcoholic fatty liver disease (NAFLD) activity score (NAS) and hepatoxicity, and the extent to which they are necessary or sufficient for fibrogenesis remains unaddressed. Thus, while recent findings implicate adaptive immunity in shaping hepatic inflammation, they fall short of establishing a causal role in fibrosis progression.

Here, the role of conventional T cell-mediated adaptive immunity in the fibrotic niche was interrogated by leveraging transcriptional and spatial profiling of human liver samples and genetically modified mice lacking αβ T cells (*Tcrb-/-*) in a western diet-induced model of fibrosing MASH. The findings of this study demonstrate that αβ T cells, while abundant in fibrotic tissue, are not required for fibrosis progression. γδ T cells, with either an auto-aggressive-like or IL-17A and TNFα phenotype, compensated in the absence of conventional T cells. Notably, the “core” of the fibrotic niche appears to be independent of T cells and results from a myeloid-centric program involving SPP1^+^ scar-associated macrophages (SAMs) and neutrophils that recruit CXCR6^+^ αβ and γδ T cells and promote myofibroblast activation. These results challenge prevailing assumptions about the immunological hierarchy in fibrosis and underscore the need to prioritize innate immune mechanisms as therapeutic targets in chronic liver fibrotic disease.

## Material and methods

### Mouse strains and models

All procedures performed on animals were in accordance with regulations and established guidelines, which were reviewed and approved by an Institutional Animal Care and Use Committee and an ethical review process. All experimental procedures were performed at Pfizer (Cambridge, MA). C57BL/6J (WT) and *Tcrb*-/- (strain: 002118; The Jackson Laboratory, Bar Harbor, ME) male mice, starting at 6 weeks of age, were used for chemical (carbon tetrachloride; CCl_4_) and/or diet-induced preclinical MASH (n= 6-15 per group). The mice were maintained in a thermo-neutral environment at a temperature of 30°C, humidity of 30-70%, and a 12 h-light/12 h-dark photocycle. Mice were injected with vehicle (olive oil) or CCl_4_ (Sigma; 56-23-5 2x/week at 2 µl CCl_4_/g) and/or fed a choline-deficient diet (CDA; Research Diets; A06071302) or HFD (Kcal of 40% fat, 20% fructose, and 2% cholesterol; D09100310; Research Diets, New Brunswick, NJ) for up to 40 weeks. Mice were euthanized using 5% CO2. Control animals were maintained on normal chow diet.

### Human tissue sections

Human paraffin-embedded (FFPE) tissue sections were obtained anonymized from BioIVT with written informed consent. An institutional review board or independent ethics committee reviewed and approved relevant study documents, including the protocol. Study conduct adhered to the ethical guidelines of the 1975 Declaration of Helsinki principles, the Council for International Organizations of Medical Sciences international ethical guidelines, and all applicable laws and regulations. Human liver tissue samples sourced from vendors followed protocols that were reviewed and approved by appropriate regulatory and ethics committees. Patient metadata is available in Table S1.

### Tissue processing

Livers from mice were collected on ice in cold RPMI 1640 medium, minced with a razor blade, and collagenase-digested (100 U/ml; Sigma-Aldrich), shaking at 37°C for 1 h. After digestion, liver tissues were filtered through 70 μm nylon mesh. Hepatocytes were removed by slow centrifugation (50 g for 5 min) and non-parenchymal cells were enriched by centrifugation through a 25% Percoll (GE Life Sciences) gradient at 850 g for 15 min. Erythrocytes were lysed with ammonium-chloride-potassium buffer (Thermo Fisher Scientific). Cells were subsequently distributed either for flow cytometry, restimulation with PMA/Ionomycin, or scRNA-seq.

### Multiplex cytokine analysis

Intrahepatic leukocytes (IHLs) were resuspended at a concentration of 5 million cells per ml in RPMI with 10% fetal bovine serum (FBS) and 1× stimulation cocktail (eBioscience) for 18 h. Supernatants were collected and stored at -80°C. Multiplex cytokine/chemokine quantification was performed using 25 μl of supernatant post-stimulation (LEGENDplex; BioLegend). Analysis was performed using BioLegend Cloud analysis software. Data were expressed as fold change normalized to control, chow-fed mice.

### Flow cytometry

Mouse and human cells were washed with phosphate-buffered saline (PBS) and stained for 20 min at room temperature with LIVE/DEAD fixable aqua viability dye (Invitrogen). Cells were washed with fluorescence-activated cell sorting (FACS) buffer and subsequently fixed with Foxp3/Transcription Factor Fixation/Permeabilization buffer (Invitrogen). After washing (PermWash Buffer; Invitrogen), cells were resuspended in freezing solution (90% FACS buffer/10% dimethyl sulfoxide (DMSO)) and stored at -80°C. Once thawed, cells were washed, and Fc receptors were blocked with TruStain FcX (mouse or human, as appropriate; BioLegend) before staining with antibodies from Table S2. Cells were then washed, filtered through 60 μm mesh, and run on a BD FACSymphony flow cytometer (BD). Data were analyzed with Omiq.ai, using a DownSample and UMAP (Uniform Manifold Approximation and Projection) workflow.

### cycIF staining and analysis

Human and mouse cyclic immunofluorescence (cycIF) was performed as described^22,23^, using antibodies listed in Tables S3 or S4, respectively . Briefly, human or mouse liver FFPE sections were deparaffinized, rehydrated, and subjected to heat-induced (Citrate PH 6) epitope antigen retrieval. Slides were stained at 4°C overnight with primary antibodies in blocking buffer solution (PBS, 0.05 % Tween-20, 5% bovine serum albumin, 0.1% Triton X-100). After three washes (PBS, 0.05% Tween-20), slides were stained with fluorescently conjugated donkey secondary antibodies (diluted 1:500 in blocking buffer) for 1 h at RT. Slides were again washed, DNA-stained with 4′,6-diamidino-2-phenylindole (DAPI; Invitrogen), and mounted in EverBrite Trueblack mounting medium to reduce auto-fluorescence. Acquisition was performed on an Axioscan Slide Scanner (Zeiss). Following imaging, coverslips were removed and antibodies were stripped for subsequent rounds of staining. For image processing, acquired images were first subjected to shading correction and stitching using the Zeiss ZEN blue DESK (version 3.0) image processing software. Alignment across subsequent rounds was performed on their respective nuclear (DAPI) channel in QuPath by applying the transformation to the remaining channels^24,25^. The multichannel image pyramid was generated using Bioformats2Raw and Raw2OMETiff programs (Glencoe Software, Seattle, WA) and before conversion into FCS for analyses in Omiq.ai.

### Fibrosis histology

FFPE sections were deparaffinized and incubated for 30 min in Picro Sirius Red fast green solution (Direct Red 81 0.2%, FCF 0.1% in 1.3% picric acid solution, Sigma-Aldrich). Sections were vigorously rinsed in dH_2_O, dried, mounted with Permount (Thermo Fisher Scientific), and acquired on an Axioscan Slide Scanner (Zeiss) in brightfield and with polarized light.

### TUNEL staining

Cell death was assessed using the ApopTag Peroxidase *In Situ* Apoptosis Detection Kit (Millipore-Sigma, S7101), following the manufacturer’s instructions. Briefly, mouse liver FFPE sections were deparaffinized, pretreated with proteinase K and quenched with endogenous peroxidase followed by the addition of equilibration buffer directly on each specimen. Slides were incubated with TdT enzyme in a humidified chamber at 37°C for 1 h, washed, and an anti-digoxignenin conjugate was applied to the slides kept in a humidified chamber for 30 min at room temperature. After additional PBS washes, peroxidase substrate developed the color, slides were washed, and a methyl-green counterstain was applied. Specimens were dehydrated with xylene and slides mounted with Permount. Brightfield slide images were acquired on an Axioscan Slide Scanner (Zeiss) and TUNEL-positive staining was quantified in QuPath.

### Immuno-histochemistry

FFPE slides were deparaffinized, rehydrated, and subjected to antigen retrieval as described above for cycIF. Endogenous peroxidases were quenched for 15 min at RT with 0.6% H_2_O_2_ in MeOH. Slides were washed three times (dH_2_O x2 followed by PBS x1), then incubated overnight in a humidified chamber in blocking buffer solution with anti-CD3e (Abcam, Rabbit host, cat# ab16669 and Invitrogen, Rat host, cat# MA5-16622, both 1:50) at 4°C. The following day, slides were washed with PBS three times and then incubated for 45 min at RT with biotinylated secondary antibodies (Goat anti-Rabbit IgG, Invitrogen, cat#65-6140 and Goat anti-Rat IgG, Vector Laboratories, cat#BA-9401-.5) diluted 1:300 in blocking buffer. The Vectastain Elite ABC Kit, Peroxidase (Vector Laboratories, cat#PK-6100) was applied following the manufacturer’s protocol for 30 min at RT. Slides were developed for ∼20 min using the DAB Substrate Kit Peroxidase (Vector Laboratories, cat#SK-4100). Slides were washed in dH_2_O and counterstained with Modified Mayer’s Hematoxylin (Epredia, cat#72804) for 2 min at RT. Slides were rinsed in dH_2_O and PBS sequentially and then dehydrated in 95% EtOH for 10 sec x2, 100% EtOH for 2 min x2, and xylene for 2 min x2 before mounting with Permount and drying overnight prior to brightfield imaging on an Axioscan Slide Scanner (Zeiss).

### Mouse scRNA-seq and data analyses

Cells from murine tissues were dissociated as described above, fixed in methanol following the 10X Genomics fixation protocol, and stored at -80°C (10X Genomics, Pleasanton, CA). A 3’ sequencing library was generated following protocols from 10x Genomics Single cell version 3.1. Samples from WT animals were sequenced on an Illumina NextSeq 2000. Samples from the *Tcrb*-/- animals were lipid-hashed as described^26^ and sequenced on an Illumina NextSeq 500. FASTQs were preprocessed and aligned with Cell Ranger version 8.0.1 using the mouse reference GRCm39-2024-A. Downstream processing of count matrices was performed using Seurat version 5.1.0^27^ and R version 4.4.0. Ambient RNA was corrected using SoupX version 1.6.2^28^. Samples with lipid hashing were demultiplexed according to the Seurat workflow using HTODemux^29^ with positive.quantile = 0.95. Wild-type (WT) and *Tcrb*-/- samples were merged and subjected to QC assessment using the Add_Cell_QC_Metrics functionality from scCustomize version 3.0.1 (RRID:SCR_024675, https://zenodo.org/records/17575311). Following uniform filtration, samples were split into WT and *Tcrb*-/- groups and analyzed in parallel following the Seurat workflow with the following modifiactions. Data were normalized with SCTransform with vars.to.regress = "nCount_RNA"^30^. PC utilization was determined by assessing both the number of dimensions that capture ∼90% variance and the point at which change in variation between PCs became <0.1%. Cells of interest (lymphocytes and myeloid cells) were subset from global objects, renormalized and clustered as described above, and filtered for contaminating cell types. Gene signature visualization was done with Nebulosa version 1.14^31^.

### Deconvolution with BayesPrism and immune cell signature analysis

Pre-processed scRNA-seq data of all cells from study GSE192742 were obtained from the cellxgene initiative (https://cellxgene.cziscience.com/collections/74e10dc4-cbb2-4605-a189-8a1cd8e44d8c) and pre-processed raw RNA-seq counts were obtained from GEO for study GSE135251. *CD8A* and *CD4* expression was used to identify CD8 T and CD4 T cells in the T cell cluster respectively using Seurat package^32^. To understand the cellular composition in samples from healthy donors and patients with NAFLD and NASH, we applied BayesPrism^33^ to predict the abundance of cell types by modeling gene expression levels through bulk RNA-Seq data. Input into BayesPrism included the raw counts from the bulk tissue and scRNA-seq normalized counts from GSE192742 as a reference. Predicted cell type specific expression from BayesPrism was normalized using variance stabilizing transformation using DESeq2 package^34^. Immune and Kupffer cell gene sets were scored in the raw counts from the bulk tissue using the z-score method in the hacksig package^35^. We applied linear regression to evaluate the association between abundances of cell types, counts per million (CPM) of select genes and cell signature scores with endpoints including age, histopathological stage and grade adjusting for batch and center as covariates.

### Hydroxyproline quantification

Liver and lung hydroxyproline content was quantified as previously described^11^. Tissues (100 to 200 mg) were hydrolyzed in 1 ml of 6 N HCl at 110°C overnight. Ten microliters of the hydrolyzed sample or standard were placed in 30 μl of citric acetate buffer [10 g of citric acid (5%, w/v), 2.4 ml of glacial acetic acid (1.2%, v/v), 14.48 g of sodium acetate (7.24%, w/v), and 6.8 g of sodium hydroxide (3.4%, w/v) with a total volume of 200 ml made up with sterile deionized water]. One hundred microliters of chloramine T solution (0.282 g of chloramine T, 2 ml of isopropanol, 2 ml of sterile water, and 16 ml of citrate acetate buffer) were mixed with the samples or standards and oxidized for 20 min at room temperature. Oxidized samples or standards received 100 μl of Ehrlich’s reagent (2.5 g of *p*-dimethylaminobenzaldehyde, 9.3 ml of isopropanol, and 3.9 ml of 70% perchloric acid) and incubated at 65°C for 20 min. Optical density at 550 nm was recorded and compared with the standard curve for quantification.

### Statistical analysis

Statistical tests were conducted using R v4.4.0 (https://r-project.org/) or GraphPad Prism v10 (GraphPad Software, La Jolla, CA). Testing for differential expression was performed in R using Seurat v5.1.0 and CellChat version 2.1.2. Graphs were generated in GraphPad Prism, the R packages Seurat v5.1.0^36^, ggplot2 version 3.5.1, scCustomize version 3.0.1, and Nebulosa version 1.14.0^31^ in R version 4.4.0, QuPath, or Omiq.ai. Differences were determined between two groups by Mann-Whitney test or between greater than two groups by analysis of variance (ANOVA) followed by Tukey post-hoc test.

## Results

### Adaptive and innate immune cells are enriched in the fibrotic niche of human MASH

To investigate immune cell proportions in human fibrotic MASH livers, a bulk RNA-seq dataset consisting of over 200 NAFLD, now referred to as metabolic dysfunction-associated steatotic liver disease (MASLD), patient biopsies^37^ (GSE135251) was deconvoluted to predict liver non-parenchymal cell (NPC) fraction frequencies from patients stratified by fibrosis stage (F0-F2 vs F3-F4) (Figure 1A). This analysis revealed a significant increase in the predicted frequency of CD8 T cells in patients with high Non-Alcoholic Steatohepatitis Activity Score (NAS; 6-8) or severe fibrosis (F3-F4) (Figure 1B). Immunofluorescence (IF) of human liver sections delineated α-SMA^+^ scar regions, which were enriched for CD3^+^, CD8^+^, and MHC-II^+^ cells in patients with severe (F3-F4) compared to minimal (F1-F2) fibrosis (Figure 1C). Cyclic immunofluorescence (cycIF) was performed to quantify frequencies of CD3^+^CD8^+^ T cells and MHC-II^+^ myeloid cells within liver scar regions (Figure 1D-F). Increased CD3^+^CD8^+^ T cell percentages were significantly associated with higher NAS (driven by lobular inflammation and ballooning) and fibrosis (Figure 1E,F), corroborating the bulk deconvolution results (Figure 1B). Conversely, MHC-II^+^ cell enrichment was specific to fibrosis severity and did not vary with other NAS parameters (Figure 1E,F). Collectively, these data establish that both adaptive and innate immune cells are spatially localized to the fibrotic niche in human MASH livers, though they associate with distinct disease hallmarks.

**Figure 1:**
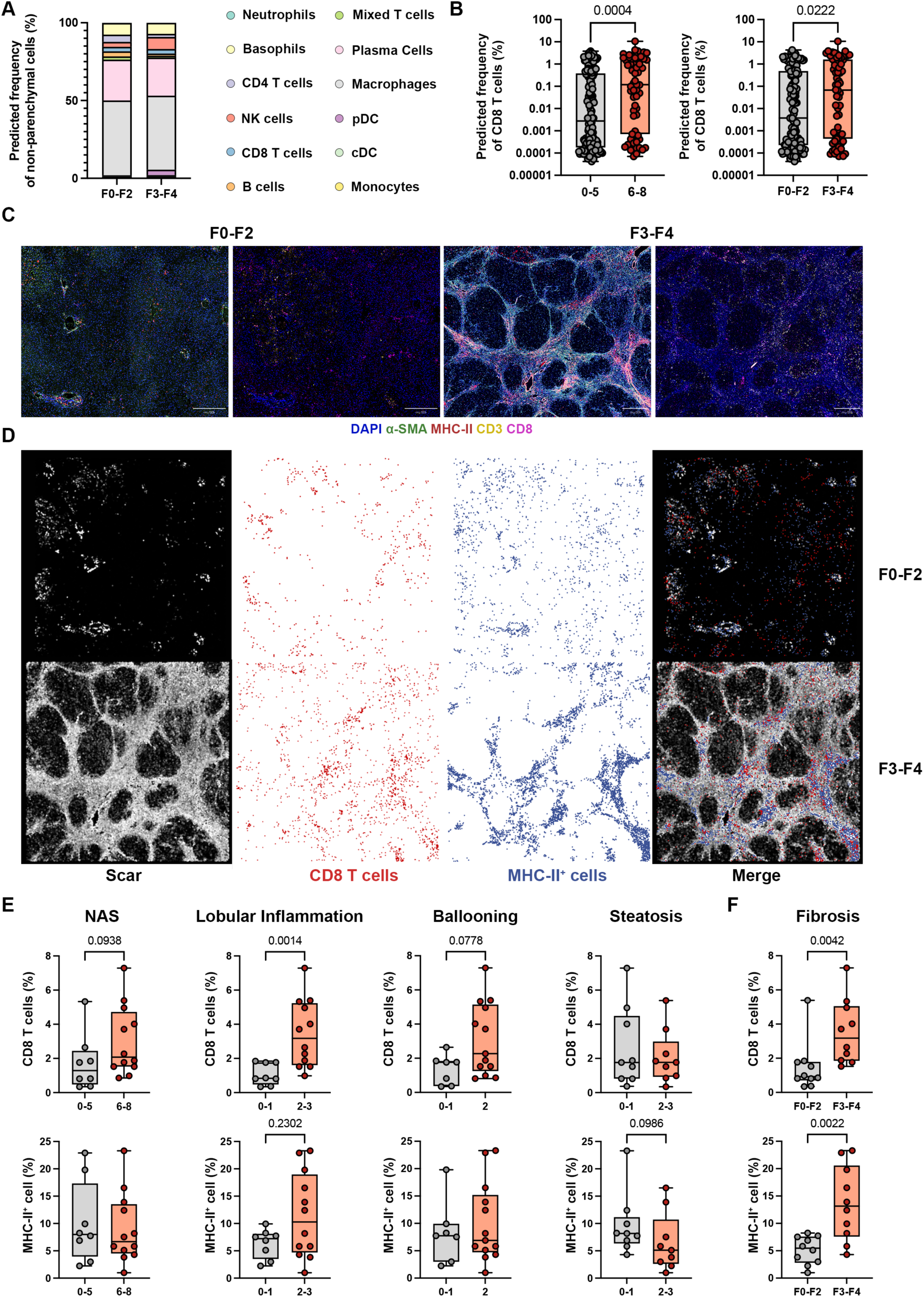
Accumulation of CD8 T cells and myeloid cells in scar in human MASH. (A) Predicted frequency from deconvoluted bulk RNA-seq (GSE135251) of immune-cells within the non-parenchymal fraction between F0-F2 and F3-F4 patients. (B) Box plot showing increased predicted frequency of CD8 T cells in NAS 6-8 and F3-F4 patients. (C) Representative immunofluorescence of α-SMA (green), MHC-II (red), CD3 (yellow) and CD8 (magenta) from F0-F2 and F3-F4 patient liver sections. (D) cycIF representative images of liver sections showing scar (α-SMA^+^ cells, grey), CD3^+^CD8^+^ T cells (red), MHC-II^+^ cells (blue) and merge overlay from patients with minimal fibrosis (F0-F2) or severe fibrosis (F3-F4). (E-F) Box plots showing CD8^+^ T cells (top) and MHC-II^+^ (bottom) cell frequencies from cycIF according to NAS, lobular inflammation, ballooning, steatosis (E) and fibrosis (F) from 20 patients with MASH. Data are presented as min – max values and median box plots. Box plot statistical analysis was performed by Mann-Whitney test.

### Auto-aggressive CD8 T cells accumulate in human fibrotic MASH and are associated with disease severity

Bulk RNA-seq datasets (GSE135251) were used to build on the established model of auto-aggressive CXCR6^+^ CD8 T cell-mediated, FASLG-dependent hepatocyte death in MASH^21^. *FAS* gene expression was significantly increased in patients with high NAS or advanced fibrosis (Figure 2A), while *FASLG*, which is expressed by NK cells and ILC1s in addition to T cells, neither correlated with NAS nor fibrosis (Figure 2B). Deconvoluted CD8 T cells, as characterized in Figure 1A, were then scored with an auto-aggressive CD8 T cell signature, as previously described^21^. CD8 T cells from patients with high NAS or advanced fibrosis were enriched for the auto-aggressive CD8 T cell signature (Figure 2C). Robust FASLG protein expression was confirmed by IF in fibrotic regions, co-localizing with CD3^+^ and CD8^+^ immune cells (Figure 2D,E). FASLG intensity was highest within CD8 T cells (Figure 2E), with the abundance of FASLG^+^ CD8 T cells increased in livers from patients with high NAS (driven by lobular inflammation) or those with advanced fibrosis (Figure 2F,G). These data suggest that auto-aggressive, FASLG-expressing CD8 T cells accumulate in the fibrotic niche in human MASH and may contribute to progressive liver injury.

**Figure 2:**
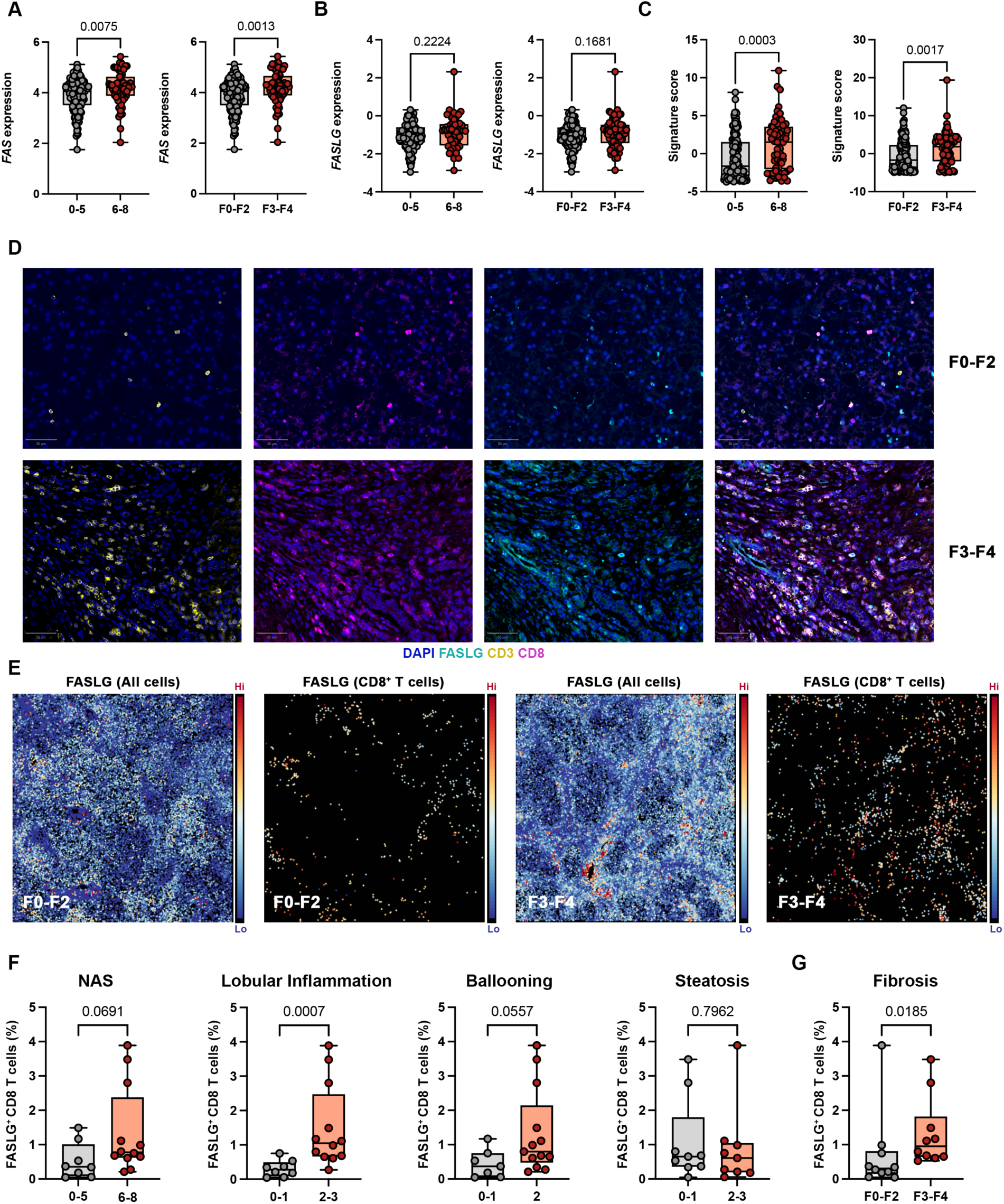
Accumulation of auto-aggressive FASLG^+^ CD8 T cells in scar in human MASH. (A-C) Bulk RNA-seq expression of *FAS* (A, global expression), *FASLG* (B, global expression) and auto-aggressive signature score (C, in deconvoluted CD8 T cells) based on fibrosis score (F0-F2, grey vs F3-F4, red). (D) Representative IF images of CD3 (yellow), CD8 (magenta) and FASLG (cyan) from F0-F2 and F3-F4 MASH livers. (E) cycIF representative images showing expression intensity (blue=low and red=high) of FASLG in all cells vs CD8 T cells from F0-F2 and F3-F4 MASH livers. (F-G) Box plots showing the frequency of FASLG^+^ CD8 T cells with NAS, lobular inflammation, ballooning, steatosis (F) and fibrosis (G). Data are presented as min – max values and median box plots. Box plot statistical analysis was performed by Mann-Whitney test.

### CD8 T cells preferentially accumulate in preclinical models of fibrosing MASH

The presence of T cells was further characterized across preclinical MASH and fibrosis models induced in *C57BL/6* wild-type (WT) mice at 6 weeks of age by chemical compounds (*i.e.,* CCl_4_) and/or dietary regimens (*i.e.,* choline-deficient (CDA) or HFD consisting of 40% fat, 20% fructose, 2% cholesterol). Fibrosis was present in all mice, as assessed by picro Sirius Red staining and imaged by either brightfield or polarized light, the latter providing improved and specific visualization of intercellular fibrosis (Figure 3A). Lymphocytes, identified by CD3 IHC staining, accumulated around the “crown like” structures of fatty hepatocytes (Figure 3B), consistent with the reported auto-aggressive T cell function of killing lipid-laden hepatocytes in proximity^21^. The proportion of CD8 T cells was significantly increased only in MASH livers with significant steatosis, such as CDA and HFD, but not CCl_4_ alone (Figure 3C). Thus, the 40-week HFD model, mimicking MASH with fibrosis and enrichment of CD8 T cells, was selected as the optimal model for this study^38^. Single-cell RNA-seq (scRNA-seq) profiling of chow and HFD livers at 40 weeks revealed a diverse immune landscape (Figure 3D, Supplementary Figure 1A,B). Livers of HFD-fed mice exhibited marked increases in the frequencies of CD8 T cells and myeloid cells (*i.e.,* monocytes/macrophages, dendritic cells), reflecting an exacerbated inflammatory state (Figure 3E), similar to human disease.

**Figure 3.**
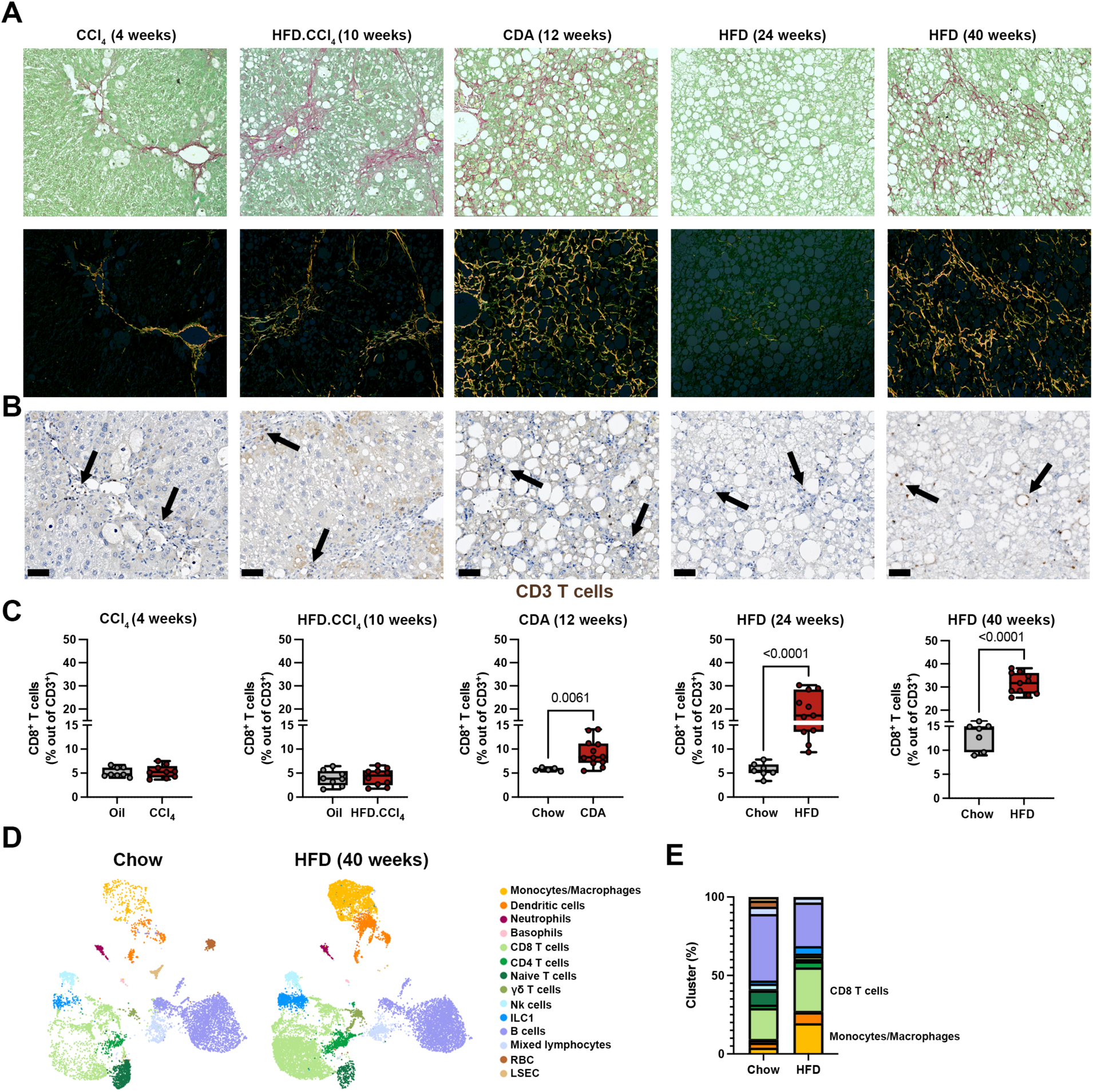
Chronic high-fat diet (HFD) leads to increased frequency of CD8 T cells in preclinical MASH models. (A) Representative brightfield (top) and polarized light (bottom) picro Sirius Red images from 4 weeks of CCl_4_, 10 weeks of HFD/CCl_4_ ,12 weeks of CDA, or 24 and 40 weeks of HFD, respectively. (B) Representative immunohistochemistry images of CD3 T cells in hepatic fibrotic regions of mice challenged with CCl_4_, HFD/CCl_4_, CDA, 24 or 40 weeks HFD, respectively. (C) Box plots showing the frequency of intrahepatic CD8 T cells from mice challenged with CCl_4_, HFD/CCl_4_, CDA, 24- or 40 weeks HFD measured by flow cytometry, respectively. (D) UMAP visualization of immune cells from scRNA-seq from chow (left) and HFD (right) livers at 40 weeks post study start (n=3 mice per group). (E) Stacked bar chart showing the relative frequency of immune cells from scRNA-seq. Data are presented as min – max values and median box plots. Box plot statistical analysis was performed by Mann-Whitney test.

### CD8 T cells display distinct transcriptional and phenotypic features linked with auto-aggression and activation in livers of chronic HFD-fed mice

To further characterize adaptive inflammation in HFD-fed mice, lymphocytes from scRNA-seq were subclustered and annotated (Figure 4A, Supplementary Figure 2A). Remodeling of the lymphocyte landscape in livers of HFD-fed mice was evident by the increase of mature CD8 T cell subsets and reduction of naive and tissue-resident populations compared to chow-fed mice (Figure 4B). Auto-aggressive CD8 T cells were delineated using joint density approximation defined by a signature of key functional markers previously described (*Cxcr6*, *Il2rb*, *Pdcd1*, *Fasl*, *Cd3e*)^21^ (Figure 4C). To complement these transcriptomic observations, protein expression on CD8 T cells was profiled by UMAP dimensionality reduction and unbiased clustering of flow cytometry data (Figure 4D, E). Ten phenotypically distinct CD8 T clusters were identified (C1-C10) (Figure 4D), with clusters C1, C6-C10 predominantly enriched in livers of HFD-fed mice (Figure 4D, E). Cluster C1 presented high expression of auto-aggressive and exhaustive CD8 T cell markers, including FASLG and PD1, respectively (Figure 4F). Chronic HFD increased the frequency of CD44^+^FASLG^+^CXCR6^hi^ CD8 T cells, reduced homeostatic tissue-resident macrophage- (TRM)-like cells (CD69^+^PD1^+^), and drove effector populations toward an exhausted phenotype (CD69^-^PD1^+^) (Figure 4G-J).

**Figure 4.**
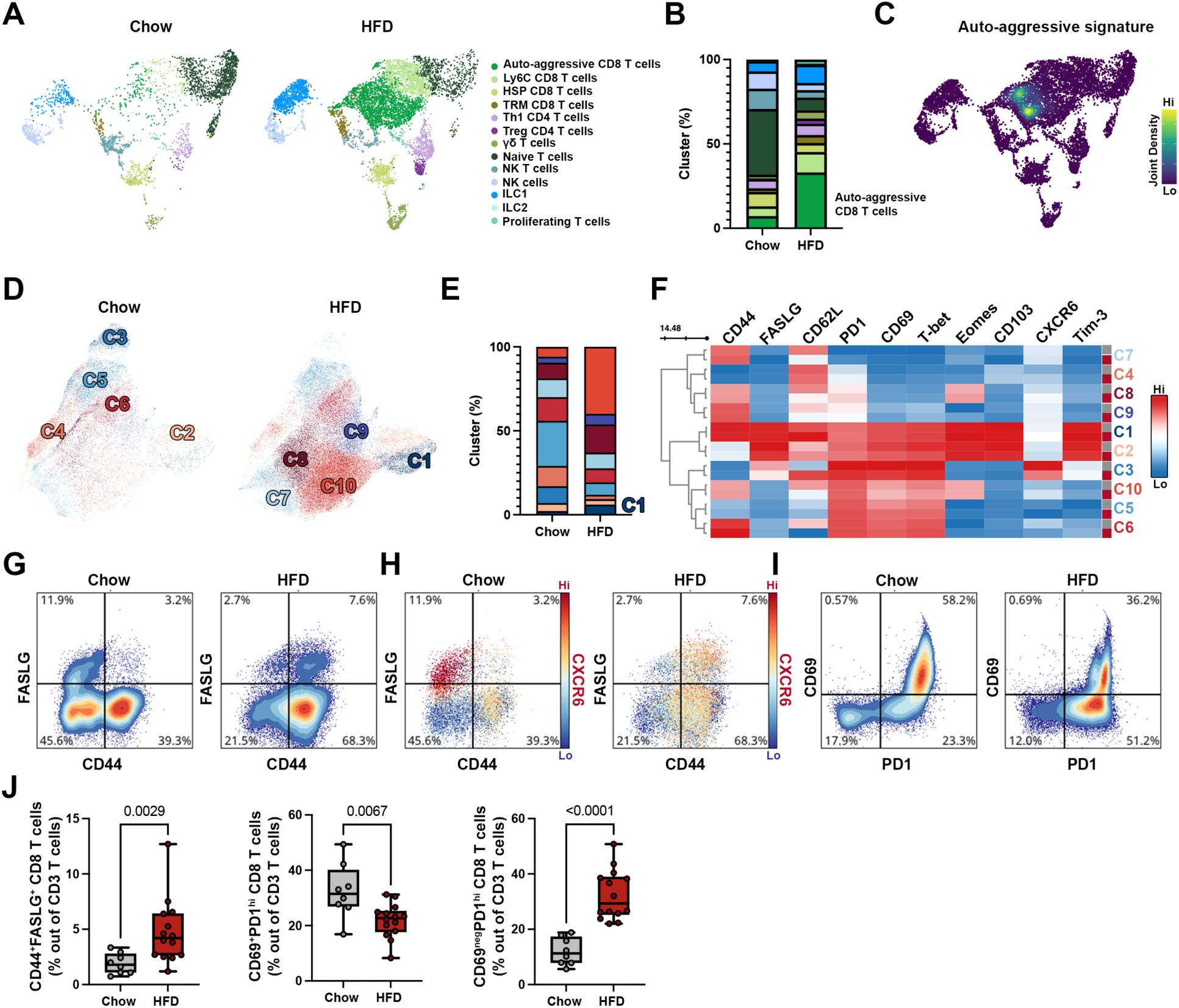
**Accumulation of FASLG^+^ auto-aggressive CD8 T cells in chronic HFD-fed mice.** (A) UMAP visualization of lymphocytes from chow (left) and HFD (right) livers after 40 weeks. (B) Stacked bar chart showing the frequency of different lymphocyte subsets from panel A and the accumulation of auto-aggressive CD8 T cells (green) in HFD livers. (C) Joint density approximation of the auto-aggressive signature (*Cxcr6*, *Il2rb*, *Pdcd1*, *Fasl*, *Cd3e*) enabling the identification of FASLG^+^ auto-aggressive CD8 T cells. (D) UMAP visualization of 10 CD8 T cell clusters identified by flow cytometry. (E) Stacked bar chart showing relative frequency of the 10 CD8 T cell clusters, shown as C1 to C10 from bottom to top, between chow and HFD livers. (F) Clustered heatmap showing the relative log2FC change in the MFI of markers on the 10 CD8 T cell clusters (grey=chow and red=HFD). (G-H) Representative FACS plots showing increased frequency of CD44^+^FASLG^+^ CD8 T cells (G), overlayed with CXCR6 MFI (H). (I) Representative FACS plots showing the decreased frequency of tissue resident memory-like CD8 T cells (CD69^+^PD1^+^) in HFD livers. (J) Box plots showing the relative frequency of CD44^+^FASLG^+^CXCR6^+^ (auto-aggressive) CD8 T cells (left), CD69^+^PD1^hi^ CD8 T cells (middle) and CD69^neg^PD1^hi^ CD8 T cells (right). Data are presented as min – max values and median box plots. Box plot statistical analysis was performed by Mann-Whitney test.

### Livers of HFD-fed *Tcrb-*deficient mice show persistent cell death and scarring with rebound γδ T cell accumulation

To assess the therapeutic potential of autoimmunity disruption in chronic diet-induced liver injury and fibrosis, *Tcrb-*deficient mice (*Tcrb*-/-), which lack conventional TCRαβ T cells, were challenged with the 40-week HFD. All HFD-fed mice exhibited progressive weight gain, with *Tcrb-*deficient mice showing a modest but significant reduction in body weight increase compared to WT counterparts (Figure 5A,B). Liver index was comparable between genotypes and remained markedly elevated relative to chow-fed baseline (dotted line, Figure 5C). Serum transaminases (*i.e.,* ALT and AST) were lower in HFD-fed *Tcrb-*deficient compared to WT mice (∼450 and ∼600 ALT U/L, respectively), but remained higher than chow-fed baseline levels (∼50 ALT U/L), indicating persistent hepatocellular injury (Figure 5D,E). Histological assessment revealed extensive steatosis and collagen deposition in both WT and *Tcrb*-deficient mice fed HFD, with no significant difference in fibrosis (Figure 5F,I). Although liver injury was reduced, cell death assessed by TUNEL^+^ staining was increased in *Tcrb*-deficient mice (Figure 5G,J), while cleaved Caspase-3 showed no significant change (Figure 5H,K), contrasting with previous reports in mice lacking both conventional αβ and γδ T cells (*Tcrb/d*-/-)^21^. As expected, *Tcrb*-deficient mice lacked conventional CD8 T cells; however, this was accompanied by a significant rebound or compensatory accumulation of intrahepatic γδ T cells (Figure 5L). These data demonstrate that the loss of TCRαβ cells did not mitigate chronic HFD-induced cell death or fibrosis, but rather promoted alternative and potentially compensatory T cell responses.

**Figure 5.**
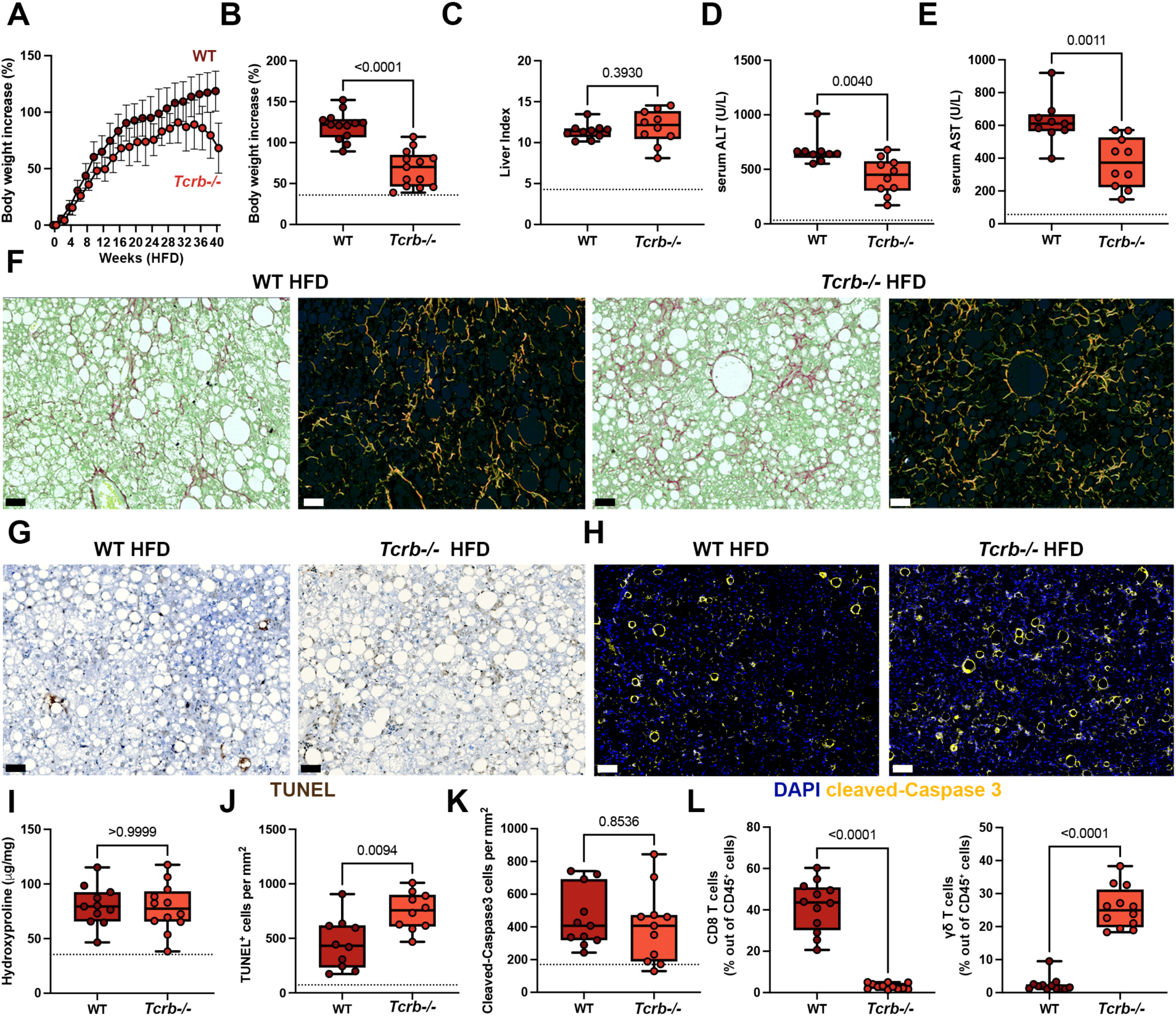
*Tcrb* deficiency does not mitigates steatosis, epithelial/hepatocellular death or fibrosis in chronic HFD-fed mice. (A-E) Body weight curves (A), percent increase in body weight after 40 weeks of HFD (B), liver index (C), serum ALT (D) and serum AST (E) between WT (dark red) and *Tcrb-/-* (red) HFD-treated mice. (F-H) Representative picro Sirius Red brightfield and polarized light (F), TUNEL staining (G) and cleaved-Caspase 3 immunofluorescent (H) images from WT and *Tcrb-/-* HFD-treated mice. (I-J) Box plots showing the quantification of hydroxyproline (I), TUNEL^+^ cells per mm^2^ (J) and cleaved-Caspase 3^+^ cells (K) between WT and *Tcrb-/-* HFD-treated mice. (L) Box plots showing the relative frequency of conventional CD8 T cells and γδ T cells between WT and *Tcrb-/-* HFD-treated mice. Data are presented as min – max values and median box plots. Box plot statistical analysis was performed by Mann-Whitney test.

### HFD-fed *Tcrb-*deficient mice present exacerbated hepatic pro-fibrotic type 3 inflammation

Liver samples from *Tcrb*-deficient mice fed control chow or HFD were also profiled by scRNA-seq (Supplementary Figure 3A-D) and intrahepatic lymphocytes were subclustered, revealing a diverse γδ T cell compartment (Figure 6A, Supplementary Figure 3E). Notably, application of the functional auto-aggressive signature (*Cxcr6*, *Il2rb*, *Pdcd1*, *Fasl*, *Cd3e*) identified a population of γδ T cells with transcriptional similarity (Figure 6B). Furthermore, a cluster with co-localized expression of type 3 cytokines, namely *Il17a* and *Tnf,* was highly prevalent during chronic HFD (Figure 6A-C). Intrahepatic leukocytes (IHLs) were then isolated and restimulated from WT and *Tcrb*-deficient livers for functional assessment of cytokine output (Figure 6D). Compared to WT, HFD-fed *Tcrb*-deficient mice released increased levels of type 1 pro-inflammatory (IL6, TNFα) and type 3 pro-fibrotic (IL17A, GMCSF) factors, alongside T cell-inducing cytokines (IL-12p40, IL-12p70, IL-23a), but not type 2 cytokines (IL-5) (Figure 6D,E). Particularly, IL6, IL-17A and TNFα were significantly elevated in IHLs from HFD-fed *Tcrb*-deficient mice (∼550, ∼1000 and ∼600 pg/mL, respectively) when compared to their WT counterparts (∼175, ∼600 and ∼100 pg/mL, respectively) (Figure 6E). These findings suggest that the absence of conventional CD8 T cells promotes compensatory γδ T cell expansion in the liver and a pro-fibrotic cytokine milieu dominated by IL-17A and TNFα, which may synergize to drive hepatotoxicity and fibrotic remodeling.

**Figure 6.**
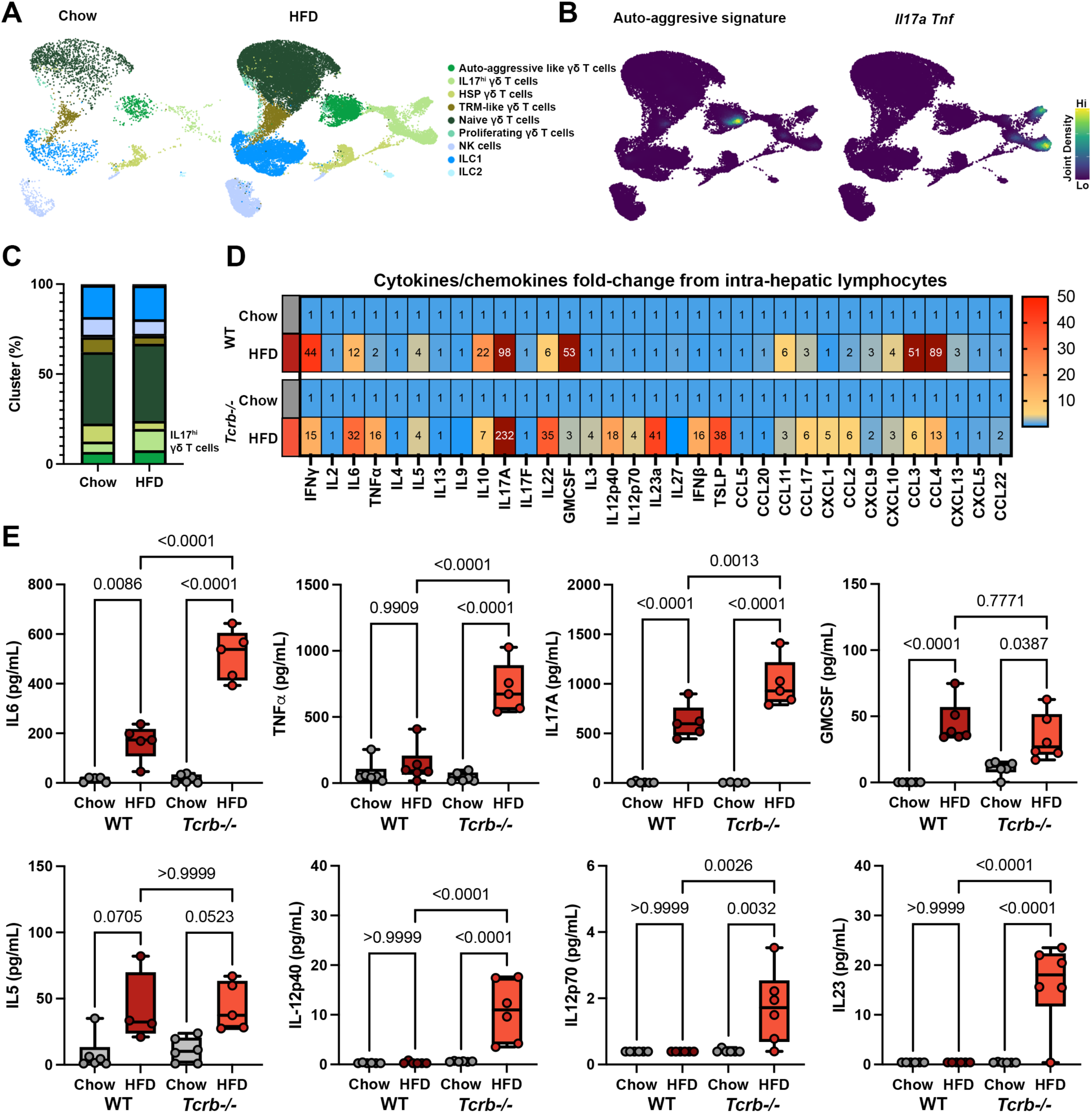
Loss of conventional CD8 T cells exacerbates pro-fibrotic type 3 inflammation. (A) UMAP visualization of lymphocytes from livers of *Tcrb-/-* mice on chow or HFD at 40 weeks by scRNA-seq. (B) Joint density expression of the auto-aggressive signature (*Cxcr6*, *Il2rb*, *Pdcd1*, *Fasl*, *Cd3e*) (left), *Il17a* and *Tnf* (right). (C) Stacked bar chart showing the frequency of lymphocytes from *Tcrb-/-* mice on chow or HFD for 40 weeks. (D) Heatmap of cytokines, fold change to chow (grey) mice, from re-stimulated IHLs isolated from WT (dark red) and *Tcrb-/-* (red) HFD mice after 40 weeks. (E) Box plots showing the intrahepatic levels of type 3 (IL-6, TNFα, IL17A, GMCSF), type 2 (IL-5) and T-cell inducing cytokines (IL-12p40, IL-12p70, IL-23) between chow-fed (grey) and chronic HFD-fed WT (dark red) and *Tcrb-/-* (red) mice. Data are presented as min – max values and median box plots. Box plot statistical analysis was performed by ANOVA and Tukey post-hoc test.

### Pro-fibrotic SPP1^+^ scar-associated macrophages (SAMs) persist in MASH livers in the absence of conventional αβ T cells

Given that *Tcrb*-deficiency failed to prevent HFD-induced fibrosis in mice, myeloid subsets were subsequently investigated. Livers of HFD-fed mice analyzed by scRNA-seq revealed an increase in myeloid diversity in HFD-fed animals compared to chow-fed controls of both WT and *Tcrb*-deficient mice (Figure 7A-F, Supplementary Figure 4A,B). In both genotypes, HFD increased the abundance of SPP1^+^ SAMs, defined by the expression of the Fab5 signature (*Cd9*, *Trem2*, *Cd63*, *Fabp5*, *Gpnmb*, *Spp1*)^23^ (Figure 7B,E). The presence of SPP1^+^ SAMs was assessed at the protein level by flow cytometry, where HFD led to the specific accumulation of CD9^+^TREM2^+^Galectin-3^+^GPNMB^+^ cells within F4/80^+^Ly6C^lo^ macrophages, in both WT and *Tcrb*-deficient mice (Figure 7G). While the relative abundance of myeloid cells was consistent across conditions, HFD led to a significant increase in CD9⁺TREM2⁺Galectin-3⁺GPNMB⁺ macrophages compared to normal chow, irrespective of genotype (Figure 7H). Heatmap profiling of flow cytometry-based protein expression further showed the upregulation of SAM-specific markers in fibrotic livers of HFD-fed mice, with comparable expression patterns between WT and *Tcrb*-deficient mice (Figure 7I). Finally, cycIF of WT and *Tcrb*-deficient HFD livers showed a fibrotic niche architecture characterized by the accumulation of GMCSF^+^IL17A^+^MMP9^+^ neutrophils, SPP1^+^GPNMB^+^FABP5^+^ SAMs^23^, and RBPJ^+^ macrophages^39^ in proximity of Desmin^+^ myofibroblasts forming crown-like structures (Figure 7J). Thus, HFD-induced SPP1^+^ SAM expansion in liver disease occurs independently of conventional αβ T cells, implicating macrophage-driven mechanisms in fibrosis persistence.

**Figure 7.**
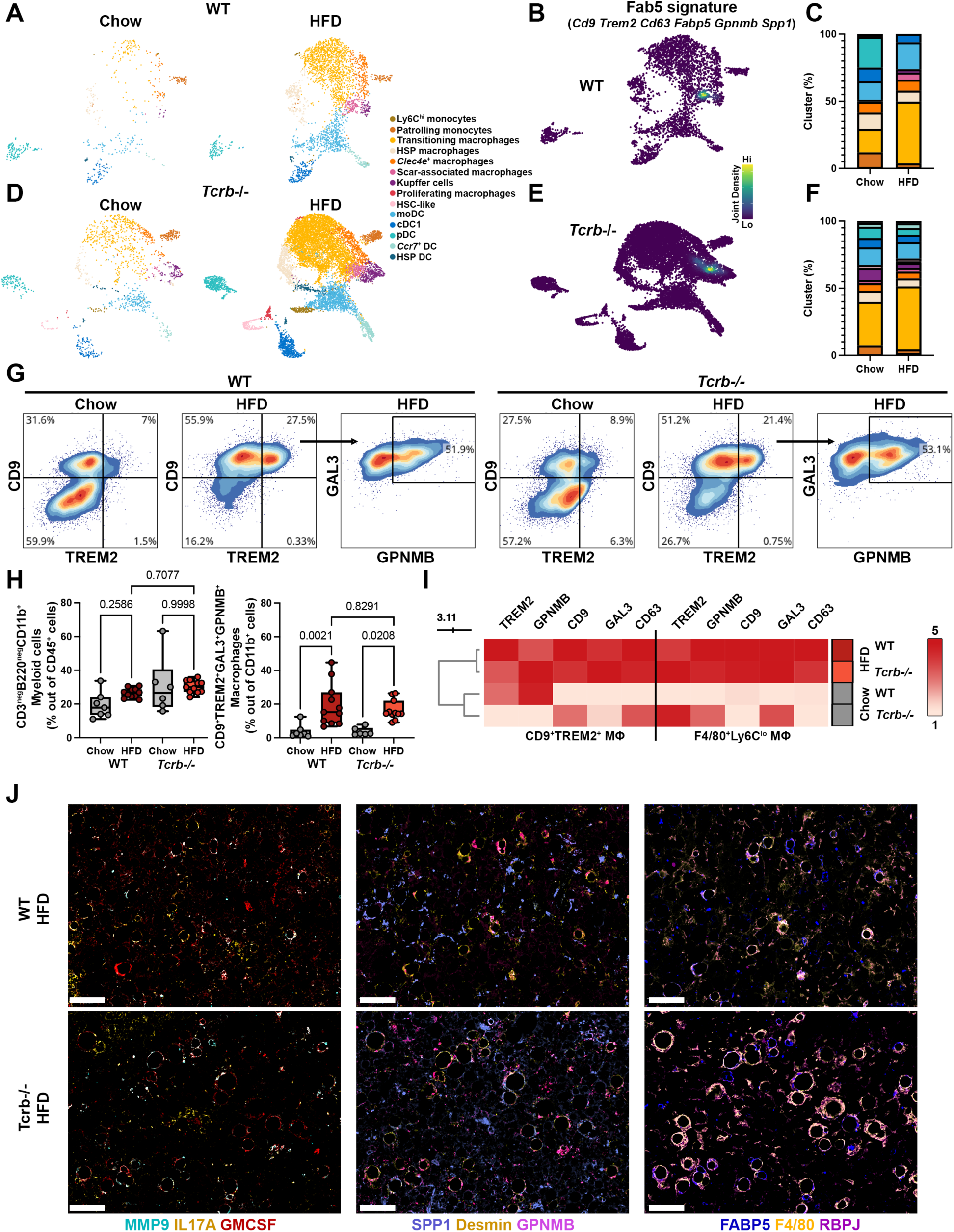
**SPP1^+^ scar-associated macrophages (SAMs) accumulate and persist in fibrosis, independently of conventional CD8 T cells.** (A,D) UMAP visualization of myeloid cells from livers of WT (A) and *Tcrb-/-* (D) mice after 40 weeks of chow (left) or HFD (right) by scRNA-seq. (B,E) Joint density expression of the pro-fibrotic Fab5 signature (*Spp1, Gpnmb, Fabp5, Cd63, Cd9*) in WT (B) and *Tcrb-/-* (E) mice. (C,F) Stacked bar chart showing the frequency of myeloid cell clusters between chow and HFD WT (C) and *Tcrb-/-* (F) mice after 40 weeks. (G) Representative FACS plots showing the identification of SPP1^+^ SAMs by flow cytometry using co-expression of CD9, TREM2, Galectin-3 and GPNMB in F4/80^+^Ly6C^lo^ macrophages from WT (left) and *Tcrb-/-* (right) mice. (H) Box plots showing the relative frequency of myeloid cells from CD45^+^ cells (left) and the relative frequency of CD9^+^TREM2^+^Galectin-3^+^GPNMB^+^ SAMs from CD11b^+^ cells (right) between WT and *Tcrb-/-* mice after 40 weeks on either chow or HFD. (I) Clustered heatmap showing the log2FC (scaled from 1 to 5) expression of scar-associated markers on F4/80^+^Ly6C^lo^ or F4/80^+^Ly6C^lo^CD9^+^TREM2^lo^ macrophages between WT and *Tcrb-/-* mice at 40 weeks on either chow or chronic HFD. (J) Representative immunofluorescent images from WT and *Tcrb-/-* mice after 40 weeks of HFD: left panel for MMP9 (cyan), IL17A (yellow) and GMCSF (red), middle panel for SPP1 (light blue), Desmin (yellow) and GPNMB (pink) and right panel for FABP5 (blue), F4/80 (yellow) and RBPJ (magenta). Data are presented as min – max values and median box plots. Box plot statistical analysis was performed by ANOVA and Tukey post-hoc test.

### CXCL16 positive myeloid cells retain CD8 T cells in fibrotic niches of human and mouse liver

Transcriptomic cell-cell interaction analysis predicted a significant increase in the number of ligand-receptor communications between T cells and myeloid cells in livers of both WT and *Tcrb-*deficient HFD-fed mice compared to chow-fed controls (Figure 8A). Interactions were filtered to chemokine/receptor pairs upregulated in HFD and revealed CXCL16 and CXCR6 as a preserved chemotactic axis between myeloid cells and both αβ and γδ T cells within fibrotic liver tissue (Figure 8B). These findings were confirmed using human liver tissue from MASH patients with either early-stage disease (F0-F2) or advanced fibrosis (F3-F4). Increased CXCL16^+^ and GPNMB^+^ macrophages were observed by IF in α-SMA⁺ fibrotic regions (Figure 8C). CycIF analysis confirmed CXCL16^+^ localization in scar regions co-populated by CD8 T cells and SPP1^+^ SAMs (Figure 8D). Spearman correlation between the frequencies of cells identified by cycIF and components of NAS and fibrosis revealed a positive correlation between CXCL16^+^ cells and the accumulation of both total and FASLG^+^ CD8 T cells in human MASH livers (Figure 8E,F). Moreover, the frequency of SPP1⁺GPNMB⁺FABP5⁺ SAMs correlated specifically with fibrosis, over other histological features (NAS and steatosis), supporting their pro-fibrotic nature (Figure 8G,H). These findings identify CXCL16-CXCR6 signaling as a key mechanism by which SAMs and other myeloid cells orchestrate immune cell positioning, recruiting T cells within the fibrotic microenvironment. Myeloid cells, particularly through SAM expansion, contribute to bystander CD8 and γδ T cell accumulation in MASH and are thus sufficient for liver fibrosis.

**Figure 8.**
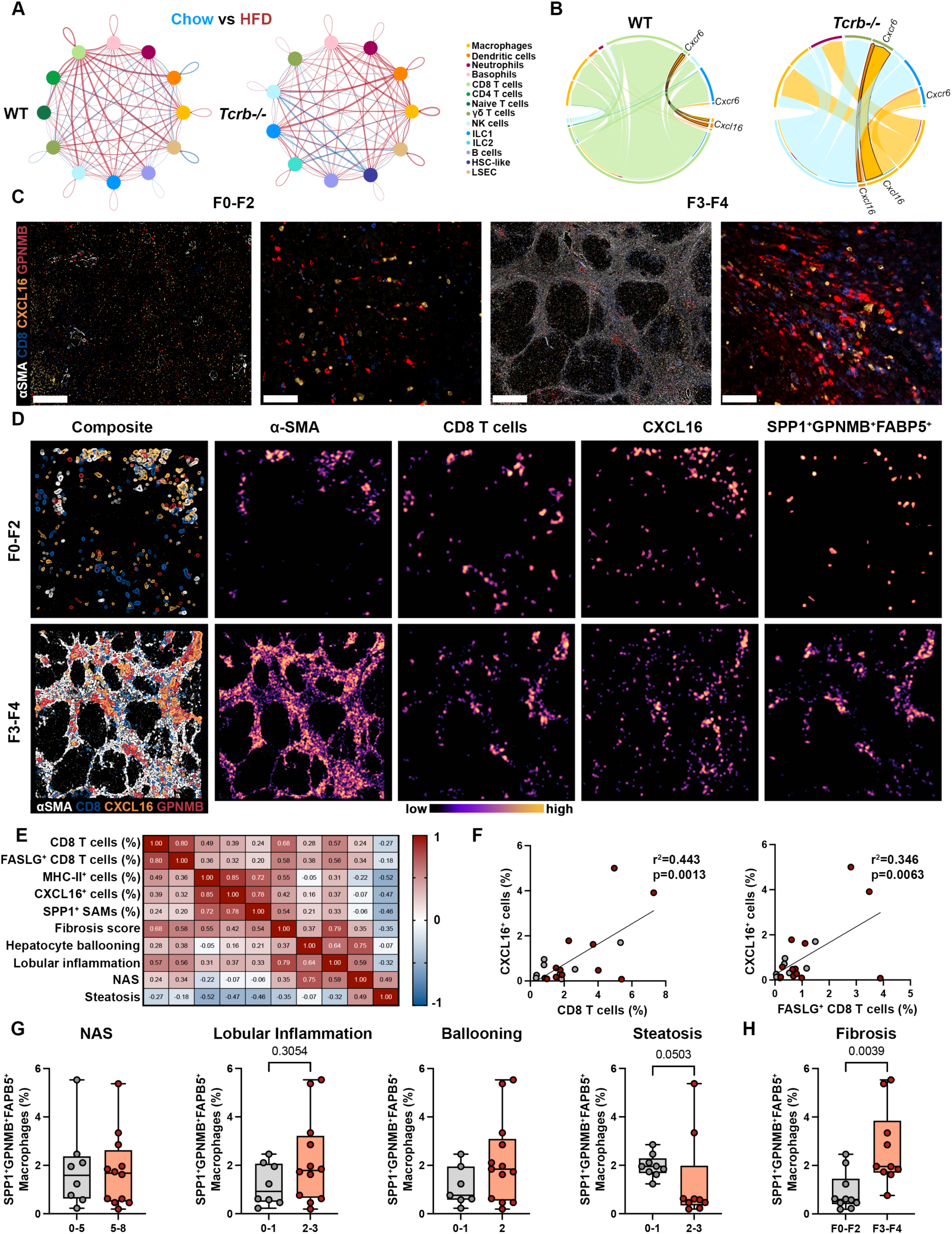
**Myeloid cell expression of CXCL16 recruits CXCR6^+^ T cells to the scar.** (A) Circle plot showing the differential number of interactions between chow (blue) and HFD (red) in WT (left) and *Tcrb-/-* (right) mice after 40 weeks by scRNA-seq. (B) Chord diagram visualizing upregulated ligands mapped to their receptors in WT (left) and *Tcrb-/-* (right) mice. Black outline highlights interactions between myeloid *Cxcl16* and conventional as well as γδ T cell *Cxcr6* relative expression by scRNA-seq. (C) Representative immunofluorescent images of α-SMA (white), CD8 (blue), CXCL16 (orange) and GPNMB (red) from F0-F2 (left) and F3-F4 (right) MASH livers. (D) cycIF representative images showing the overlaid composite density of α-SMA (white), CD8 (blue), CXCL16 (orange) and GPNMB (red) and the relative density of cells positive for α-SMA, CD8, CXCL16, and GPNMB. (E) Heatmap showing spearman correlation (-1 to 1) between the frequencies of CD8 T cells, FASLG^+^ CD8 T cells, CXCL16^+^ cells, MHC-II^+^ cells, SPP1^+^ SAMs, fibrosis, hepatocyte ballooning, lobular inflammation, NAS and steatosis. (F) Spearman correlation between the frequency of CXCL16^+^ cells and the frequency of CD8 T cells (left) or FASLG^+^ CD8 T cells (right) in MASH livers. (G-H) Box plots showing the frequency of SPP1^+^GPNMB^+^FABP5^+^ SAMs with NAS, lobular inflammation, ballooning, steatosis (G) and fibrosis (H). Data are presented as min – max values and median box plots. Box plot statistical analysis was performed by Mann-Whitney test.

## Discussion

The immunopathological landscape of MASH has often featured the accumulation of T cells, due to the metabolic syndrome, dysbiosis and hepatocellular cell death triggering chemotactic responses to the liver^4^. T cells contribute to fibrosis progression via either pro-inflammatory and pro-fibrotic cytokine secretion or cytotoxic pathways. We and others reported the dysregulation of the T effector (Th2/Th17) to T regulatory cell ratio in MASH, contributing to the pro-fibrotic type 2 and 3 mixed inflammatory milieu^11,18^. MASH livers also feature a lipotoxic environment that affects survival of CD4 T cells and leads to a significant accumulation of CD8 T cells^19^. Here, we confirmed first by deconvolution of a bulk RNA-seq dataset of over 200 patients (Figure 1A,B) and in an independent cohort of FFPE MASH livers using cycIF (Figure 1C-E) that CD8 T cell accumulation in the liver correlates with NAS, lobular inflammation and fibrosis stage. CD8 T cells may expand in livers due to the presence of XCR1^+^ cDC1s, and chronic antigenic stimulation, promoting inflammation and hepatocellular damage^20,40^. Liver homing/resident CXCR6^+^ CD8 T cells, rendered auto-aggressive through IL-15-driven transcriptional reprogramming and metabolic activation, were reported to contribute to FASLG-mediated hepatocyte apoptosis in preclinical models of MASH^21^. This auto-aggressive signature was significantly enriched in livers with high NAS (6-8) and fibrosis (F3-4), as demonstrated by deconvoluted bulk RNA-seq (Figure 2C). Using cycIF, we confirmed that auto-aggressive CD8 T cells (defined by the co-expression of FASLG, CD3 and CD8) correlated with lobular inflammation and were enriched in fibrotic scars (Figure 2D-G) in human MASH livers.

In line with previous reports, CD3 T cells were located in areas of fibrosis in both toxin- and diet-induced fibrosis models (Figure 3A, B). However, the frequency of CD8 T cells was significantly increased only in models with severe and chronic steatohepatitis such as CDA and HFD. This could explain the previous observation that adaptive immune cells may not contribute to fibrosis in toxin-dependent preclinical models. Indeed, expansion of CD8 T cells in MASH appears to be clonal^40^, as a result of lipid accumulation in the liver and potential dysbiosis. Additionally, scRNA-seq and flow cytometry confirmed that chronic western-diet (high fat, fructose and cholesterol) induced an auto-aggressive CD8 T cell phenotype (CD44^hi^CXCR6^+^FASLG^+^PD1^+^), as previously reported using CDHFD^21^. Genetic ablation of conventional αβ T cells using *Tcrb*-deficient mice led to a modest improvement of certain physiopathological features of MASH, such as percent body weight and serum ALT (Figure 5A-D). However, serum ALT remained nearly ten times higher than mice on chow diet, which was consistent with the lack of improvement in cell death, assessed by TUNEL^+^ and cleaved-Caspase-3^+^ cell counts, in livers of HFD-fed *Tcrb*-deficient mice (Figure 5G,H,J,K). Foreseeably, this was associated with no significant improvement in fibrosis, as determined by both picro Sirius Red staining and hydroxyproline quantification (Figure 5F,I). These results contrast with the recent observation linking both CD4 and CD8 T cells to the amplification of inflammation and fibrosis in MASH^41^.

The contribution of γδ T cells in the liver has been linked to both fibrosis progression and resolution, depending on their phenotype. These γδ T cells can limit hepatocellular death via the secretion of IL-22, activation of hepatic stellate cells (HSCs) via the secretion of IFNγ, and mediate clearance of activated HSCs via FAS/FASLG^42^. The role of auto-aggressive CD8 T cells in MASH was initially defined in mice lacking both αβ and γδ T cells (*Tcrb/d*-double-deficient)^21^. In our case, a significant compensatory rebound in γδ T cells in livers of *Tcrb*-deficient mice was observed after chronic HFD (Figure 5L). The γδ T cells in HFD-fed *Tcrb-/-* mice displayed a diverse array of pro-fibrotic phenotypes, including a cell cluster enriched for the functional auto-aggressive signature (*Cxcr6*, *Il2rb*, *Pdcd1*, *Fasl*, *Cd3e*), originally described in CD8 T cells. Another cluster of γδ T cells showed co-expression of *Il17a* and *Tnf,* consistent with the significantly higher level of IL-17A, GMCSF and TNFα in re-stimulated IHLs from *Tcrb*-deficient mice when compared to WT mice following 40 weeks of chronic HFD (Figure 6D,E). Notably, IL-17A is a pro-inflammatory and pro-fibrotic cytokine associated to fibrosis progression in many types of liver injury and tissues.

IL-17A facilitates TNF-driven hepatotoxicity in steatotic hepatocytes, amplifying Caspase-2-SREBP1/2-mediated lipogenesis and inflammation^43,44^. IL-17A also sensitizes HSCs to the action of TGFβ via stabilization of the TGF-β-RII on the surface. Altogether, our results indicate that γδ T cells expand in the absence of conventional αβ T cells and maintain a milieu favorable, skewed towards type 3 inflammation, to the development of the fibrotic niche^45^.

Fibrotic scars of both WT and *Tcrb*-deficient mice showed myeloid cell dysregulation, particularly characterized by the loss of tissue-resident Kupffer cells and accumulation of SPP1^+^ SAMs with a Fab5 signature^23^. Lack of conventional αβ T cells neither impacted the overall frequency, transcriptomic signature nor phenotype of SPP1^+^ SAMs (Figure 7A-I). Histologically, scars from WT and *Tcrb*-deficient mice showed similar accumulation of IL17A^+^GMCSF^+^MMP9^+^ neutrophils in proximity of myofibroblasts and SPP1^+^GPNMB^+^FABP5^+^ macrophages. SPP1^+^ SAMs are associated with fibrosis progression across tissues, degrade normal extracellular matrix (ECM; Collagen IV) and increase deposition of pathogenic ECM (Collagen I)^23^. Accordingly, these cells accumulated in human MASH livers with severe fibrosis at the scar border and correlated with histopathological fibrosis grade (Figure 8C,D,H). Unlike CD8 T cells, SPP1^+^ SAMs neither correlated with increased NAS, lobular inflammation, hepatocyte ballooning nor steatosis (Figure 8G), reinforcing their pro-fibrotic association. Moreover, we found increased protein expression of RBPJ (Figure 7J), a transcription factor involved in pathogenic monocyte to macrophage (MDM) differentiation, in myeloid cells associated with fibrosis in livers from HFD-fed mice^39^.

Finally, we investigated whether myeloid cells in the liver fibrotic niche may contribute to the accumulation of CXCR6^+^ T cells during MASH. Unbiased chemokine/receptor interaction predictions identified a conserved mechanism of myeloid-driven T cell recruitment to the fibrotic niche in livers of both *Tcrb*-deficient and WT HFD-fed mice (Figure 8A,B). These data show that macrophages/monocytes and dendritic cells are sources of CXCL16 that bind CXCR6 on auto-aggressive CD8 T cells and γδ T cells. Taken together, these results indicate that myeloid cells, rather than conventional adaptive immune cells, are pivotal regulators of inflammation at the fibrotic niche. While broad neutralization of myeloid cells via inhibition of CCR2 showed limited efficacy in patients, due to its broad effect, it is possible that more refined myeloid cell inhibition would be beneficial^46–50^. Preventing monocyte recruitment by CCR2 antagonism not only limits the development of pathogenic MDMs, but also the ability of monocytes to restore the pool of depleted tissue resident macrophages (*i.e.,* Kupffer cells) in MASH. As an alternative therapeutic approach, myeloid cell inhibition of RBPJ blunted differentiation towards SPP1^+^ SAMs, improved recovery of Kupffer cells and induced patrolling monocytes, reducing inflammation and fibrosis in preclinical MASH models^39^.

This study contextualizes the roles of adaptive and innate immunity within a broader immunological landscape of the fibrotic niche. Since our data indicate redundancy between CXCR6^+^ lymphocytes in MASH, we suggest that targeting the myeloid cells responsible for CXCL16 production and HSC activation in combination with anti-steatotics will yield superior anti-fibrotic activity. In conclusion, our study demonstrates that myeloid cells are master regulators of fibrosis and fibrosis-associated adaptive immunity, redefining the immune-mediated fibrotic niche in MASH.

## Abbreviations

CCl_4_: carbon tetrachloride CDA, choline-deficient diet
CDHFD: choline-deficient high-fat diet CPM, counts per million
cycIF: cyclic immunofluorescence DC, dendritic cell
DMN: dimethylnitrosamine ECM, extracellular matrix
FACS: fluorescence-activated cell sorting FBS, fetal bovine serum
FFPE: formalin-fixed paraffin-embedded
HBV and HCV: hepatitis B virus and hepatitis C virus, respectively
HFD: high-fat diet
HSC: hepatic stellate cell
IF: immunofluorescence
IHL: intrahepatic leukocyte
MASLD: metabolic dysfunction-associated steatotic liver disease
MASH: metabolic dysfunction-associated steatohepatitis
MCDD: methionine and choline-deficient diet
MDM: monocyte-derived macrophage
NAFLD: non-alcoholic fatty liver disease (*now referred to as MASLD*) NAS, NAFLD activity score
NASH: non-alcoholic steatohepatitis (*now referred to as MASH*) PBS, phosphate buffered saline
SAM: scar-associated macrophage
scRNA-seq: single cell RNA-seq
TRM: tissue-resident macrophage
WT: wild-type

## Financial support

This study was financed by Pfizer Inc.

## Conflict of interest

C.L, K.L.D., S.L., S.M.C., S.A., M.H.W., K.B., S.D., T.A.W., K.M.H., T.F. are or J.M., F.S., C.W.,

X.C., A.M.S.B. were employees of Pfizer Inc.

## Author contributions

Acquisition, analysis and interpretation of data: C.L, K.L.D., S.L., S.M.C., S.A., M.H.W., K.B, S.D.,

J.M., F.S., C.W., X.C., A.M.S.B., K.M.H., T.F. Drafting and writing of the manuscript: C.L., K.L.D,

T.F. Critical revision of the manuscript for important intellectual content: T.A.W. Study concept, design supervision: A.M.S.B., T.A.W., K.M.H., T.F.

## Data availability

The datasets generated and/or analyzed during the present study are available from the corresponding author upon reasonable, scientifically justified request.

## Acknowledgements

We thank the Pfizer Flow Cytometry, Optical Microscopy, High Content Imaging, and Next-Generation Sequencing Technology Centers. K.L.D. thanks the Pfizer Postdoctoral Fellowship Program for support.

---

**Supplementary Figure 1.**
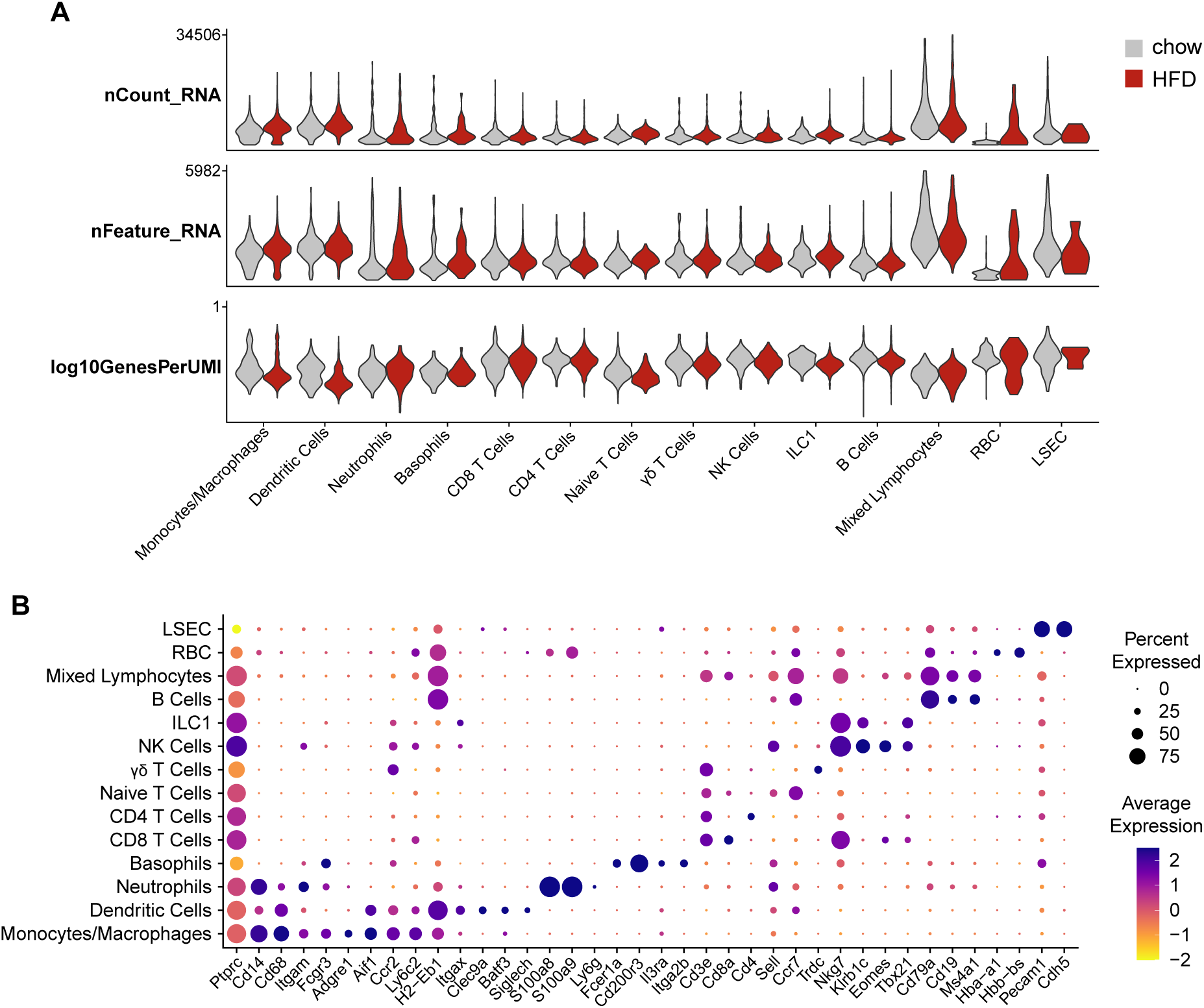
Livers of chow- and HFD-fed wild-type (WT) mice profiled by scRNA-seq. (A) Stacked violin plots of UMI, gene expression, and cell complexity by cell type in chow (grey) and HFD-fed (red) WT mice after 40 weeks of diet. (B) Dot plot of global marker gene expression in chow and HFD-fed WT mice combined.

**Supplementary Figure 2.**
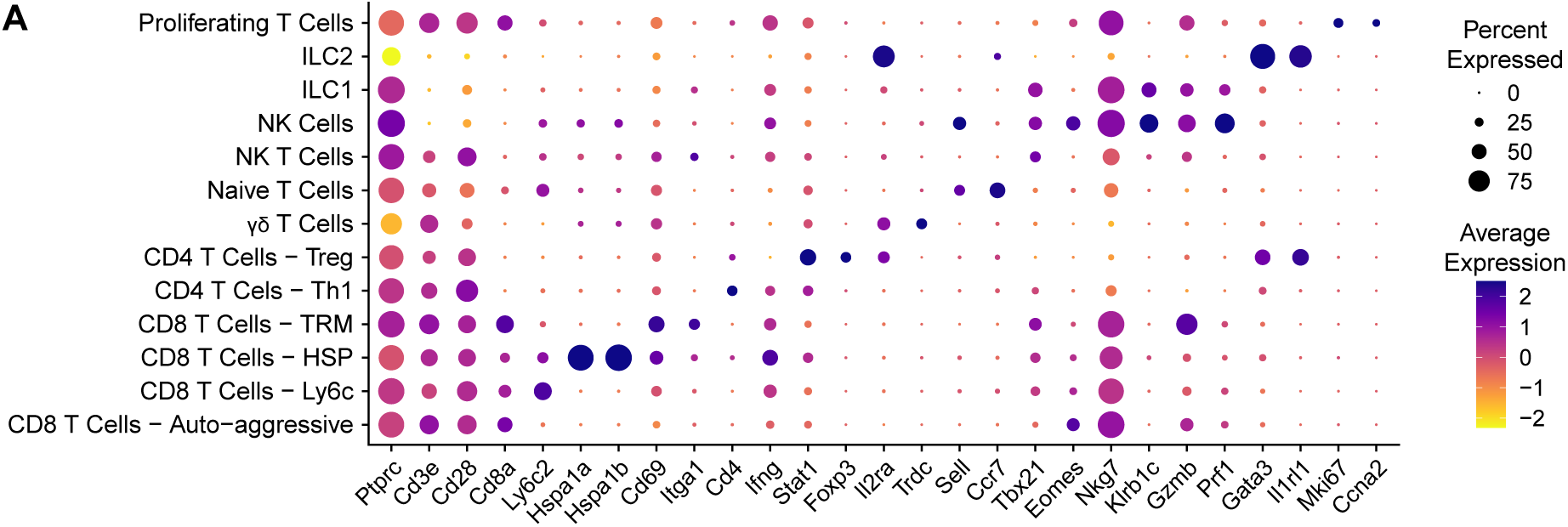
WT liver lymphocyte gene expression by scRNA-seq. (A) Dot plot of marker gene expression in subclustered lymphocytes from livers of chow and HFD-fed WT mice combined.

**Supplementary Figure 3.**
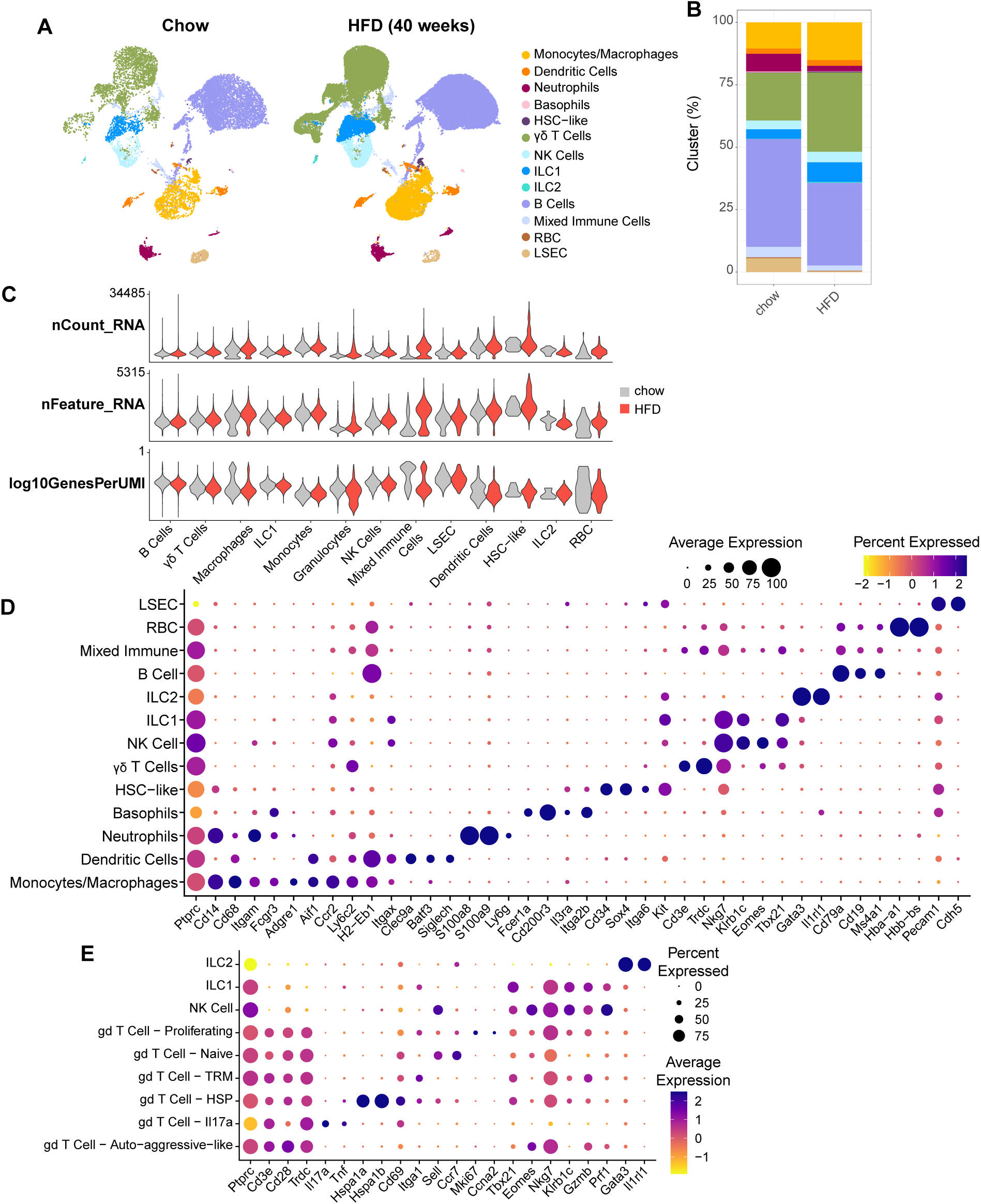
Livers of chow- and HFD-fed *Tcrb*^-/-^ mice profiled by scRNA-seq. (A) Split UMAP of scRNA-seq global cell annotations from chow and HFD-fed *Tcrb*-/- mice after 40 weeks of diet. (B) Relative abundancies of global populations detected by scRNA-seq. (C) Stacked violin plots of UMI, gene expression, and cell complexity by cell type in chow (grey) and HFD-fed (red) *Tcrb*-/- mice. (D-E) Dot plots of global (D) and lymphocyte (E) marker gene expression in livers of chow- and HFD-fed *Tcrb*-/- mice combined.

**Supplementary Figure 4.**
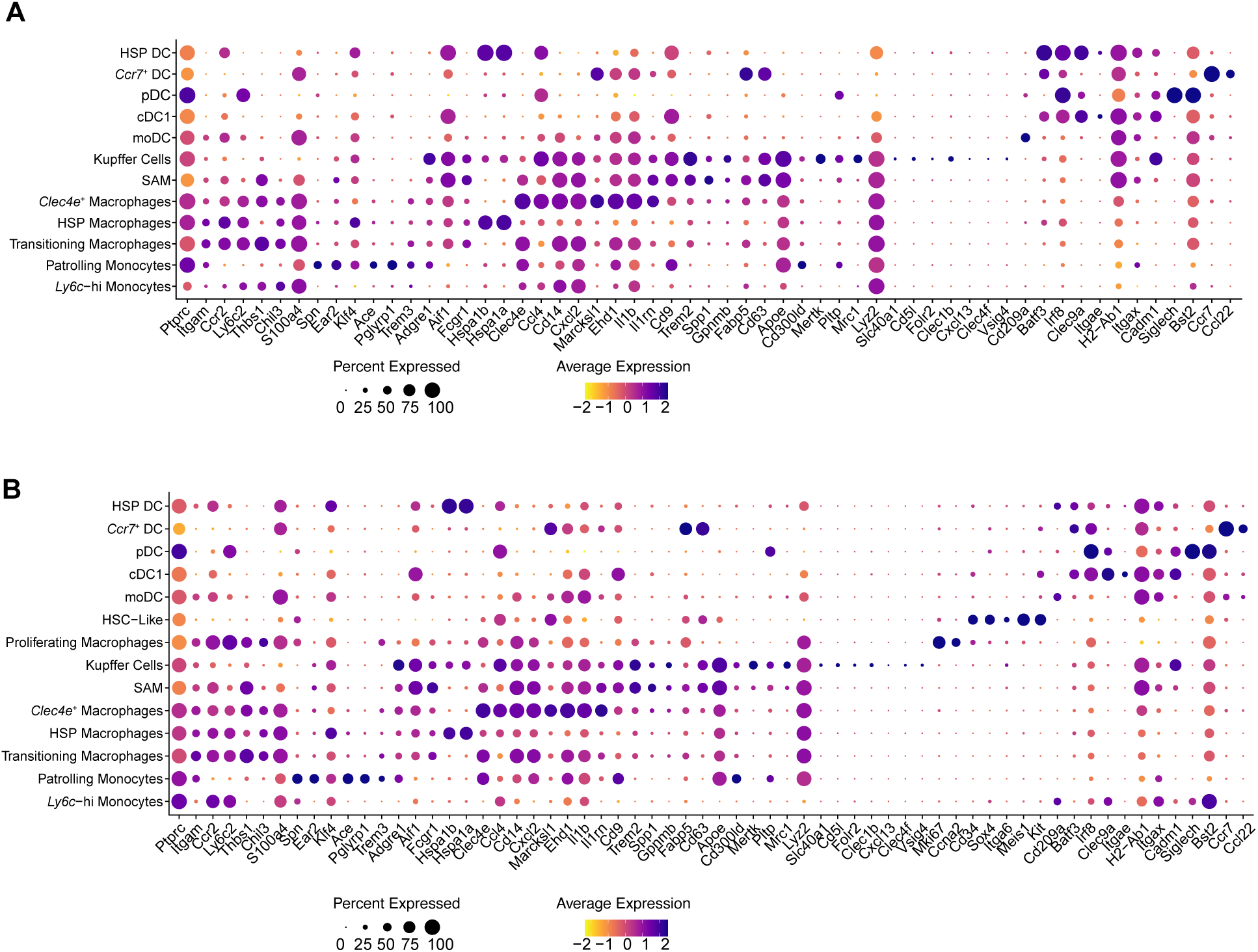
MASH liver myeloid gene expression by scRNA-seq. Dot plots of marker gene expression in combined chow- and HFD-fed WT mice (A) and *Tcrb*-/-mice (B).

**Supplementary Figure 5.**
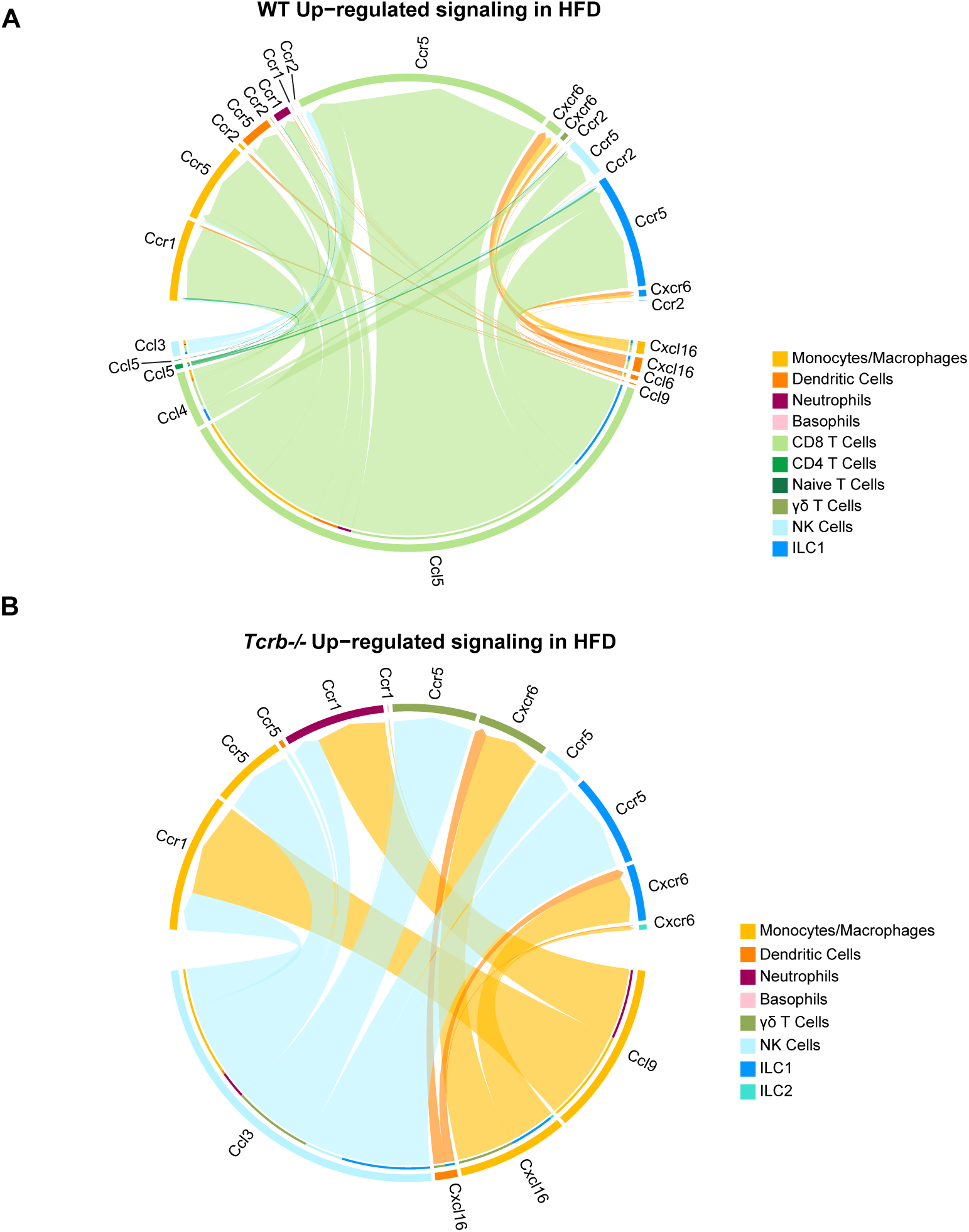
MASH liver chemokine/receptor interaction predictions from scRNA-seq. Interactions upregulated in livers of HFD-fed WT (A) and *Tcrb-/-* (B) compared to chow-fed mice.

**Table S1.** Patient metadata.

| Patient ID | Disease | Fibrosis Stage | Hepatocyte Ballooning | Lobular Inflammation | NAS | Steatosis |
| --- | --- | --- | --- | --- | --- | --- |
| F202288354388482 | MASLD/MASH | 4 | 1 | 1 | 3 | 1 |
| F269667491781714 | MASLD/MASH | 1 | 2 | 2 | 7 | 3 |
| F25230374217327 | MASLD/MASH | 3 | 0 | 3 | 3 | 0 |
| F60076066942084 | MASLD/MASH | 4 | 2 | 3 | 6 | 1 |
| F93815000985241 | MASLD/MASH | 4 | 2 | 3 | 6 | 1 |
| F164630764073724 | MASLD/MASH | 0 | 0 | 1 | 3 | 2 |
| F18870497925578 | MASLD/MASH | 3 | 1 | 2 | 3 | 0 |
| F54808383304812 | MASLD/MASH | 2 | 2 | 2 | 7 | 3 |
| F89655557507529 | MASLD/MASH | 4 | 2 | 3 | 5 | 0 |
| F160333745451172 | MASLD/MASH | 0 | 0 | 1 | 3 | 2 |
| F194072749341725 | MASLD/MASH | 4 | 2 | 3 | 8 | 3 |
| F228919843573023 | MASLD/MASH | 4 | 2 | 3 | 5 | 0 |
| F17160862857391 | MASLD/MASH | 4 | 2 | 3 | 5 | 0 |
| F51999294867643 | MASLD/MASH | 3 | 2 | 3 | 7 | 2 |
| F89045601851397 | MASLD/MASH | 2 | 2 | 3 | 7 | 2 |
| F224751406165187 | MASLD/MASH | 0 | 2 | 1 | 3 | 0 |
| F12855615291317 | MASLD/MASH | 1 | 1 | 1 | 5 | 3 |
| F47702485095320 | MASLD/MASH | 1 | 1 | 1 | 3 | 1 |
| F82540773355965 | MASLD/MASH | 0 | 1 | 1 | 5 | 3 |
| F118489267437699 | MASLD/MASH | 1 | 1 | 1 | 4 | 2 |

**Table S2.** Flow cytometry antibodies.

| Target | Fluorophore | Clone | Panel |
| --- | --- | --- | --- |
| Trem2 | FITC | 78.18 | SAMs |
| CD45 | A700 | 30-F11 | SAMs |
| GPNMB | e660 | CTSREVL | SAMs |
| CD9 | APC-Fire750 | MZ3 | SAMs |
| CD3 | BUV395 | 145-2C11 | SAMs |
| B220 | BUV395 | RA3-6B2 | SAMs |
| Siglec-F | BUV395 | E50-2440 | SAMs |
| NK1.1 | BUV395 | PK-136 | SAMs |
| CD11b | BUV787 | M1/70 | SAMs |
| CD81 | BV421 | EAT2 | SAMs |
| F4/80 | BV605 | T45-2342 | SAMs |
| Galectin-3 | PE | M3/38 | SAMs |
| Ly6G | PE-CF594 | 1A8 | SAMs |
| Ly6C | PECy7 | HK1.4 | SAMs |
| CD63 | PerCP.Cy5.5 | NVG2 | SAMs |
| Galectin-3 | PE | M3/38 | SAMs |
| Ly6G | PE-CF594 | 1A8 | SAMs |
| Ly6C | PE-Cy7 | HK1.4 | SAMs |
| F4/80 | BB515 | T45-2342 | Lineage |
| CD11c | PerCP.Cy5.5 | N418 | Lineage |
| TCRgd | BV421 | GL3 | Lineage |
| CD11b | BV605 | M1/70 | Lineage |
| B220 | BV786 | RA3-6B2 | Lineage |
| CD3 | BUV395 | 145-2C11 | Lineage |
| CD4 | BUV496 | GK1.5 | Lineage |
| CD8 | BUV787 | 53-6.7 | Lineage |
| CD68 | APC | FA-11 | Lineage |
| CD45 | A700 | 30-F11 | Lineage |
| NK1.1 | APC-Cy7 | PK-136 | Lineage |
| Siglec-F | PE | E50-2440 | Lineage |
| Ly6G | PE-CF584 | 1A8 | Lineage |
| Ly6C | PE-Cy7 | HK1.4 | Lineage |
| CD44 | A488 | KM81 | CD8 T cell |
| CD45 | A700 | 30-F11 | CD8 T cell |
| FASL | APC | MFL3 | CD8 T cell |
| CD62L | APC-Cy7 | MEL-14 | CD8 T cell |
| CD3 | BUV395 | 145-2C11 | CD8 T cell |
| CD4 | BUV496 | GK1.5 | CD8 T cell |
| CD8 | BUV787 | 53-6.7 | CD8 T cell |
| PD1 | BV421 | J43 | CD8 T cell |
| CD69 | BV605 | H1-2F3 | CD8 T cell |
| Tbet | BV785 | O4-46 | CD8 T cell |
| Eomes | PE | X4-83 | CD8 T cell |
| CD103 | PE-CF594 | M290 | CD8 T cell |
| CXCR6 | PE-Cy7 | SA051D1 | CD8 T cell |
| Tim-3 | PerCP-CY5.5 | RMT3-23 | CD8 T cell |

**Table S3.**
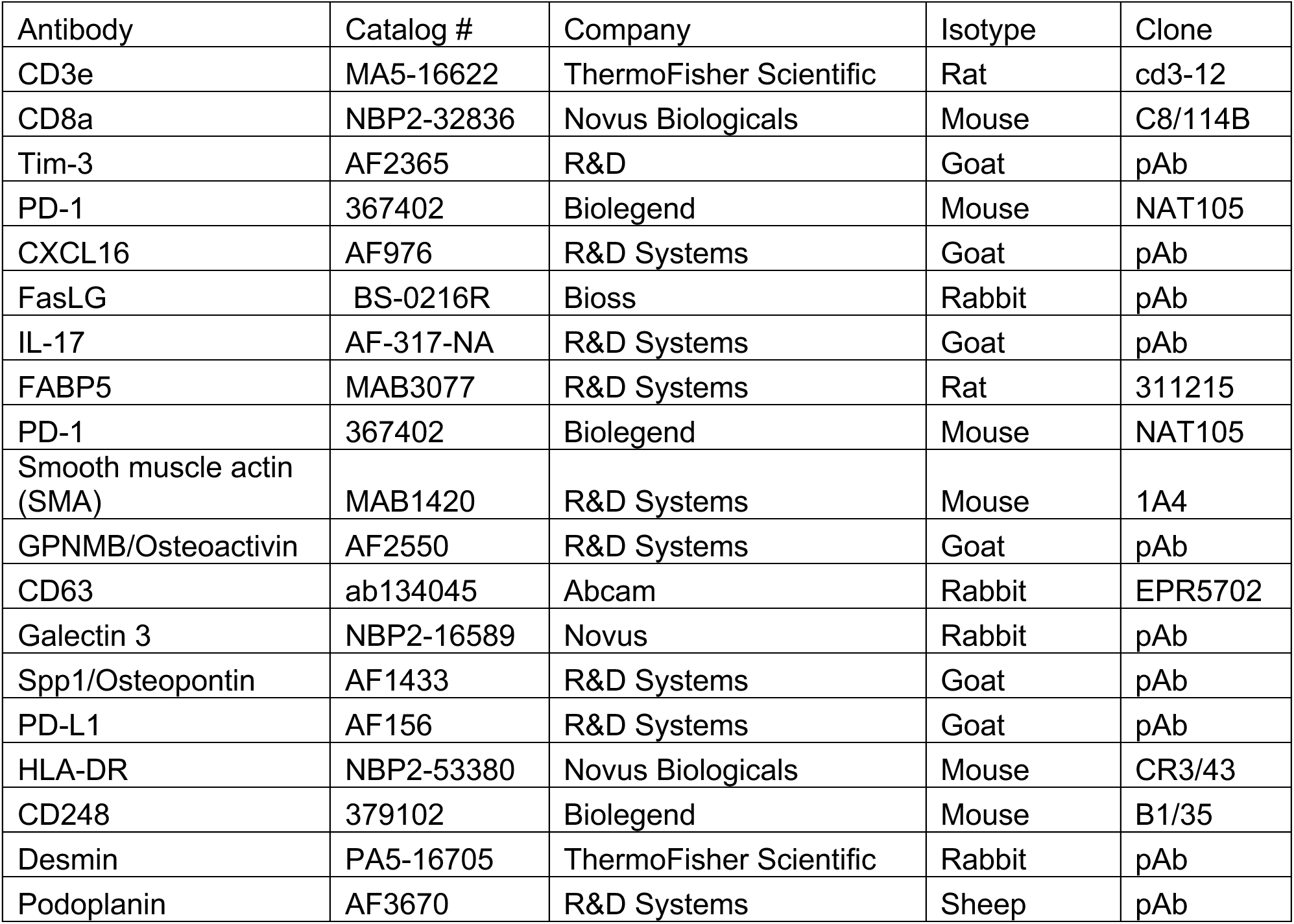
Human cycIF antibodies.

**Table S4.**
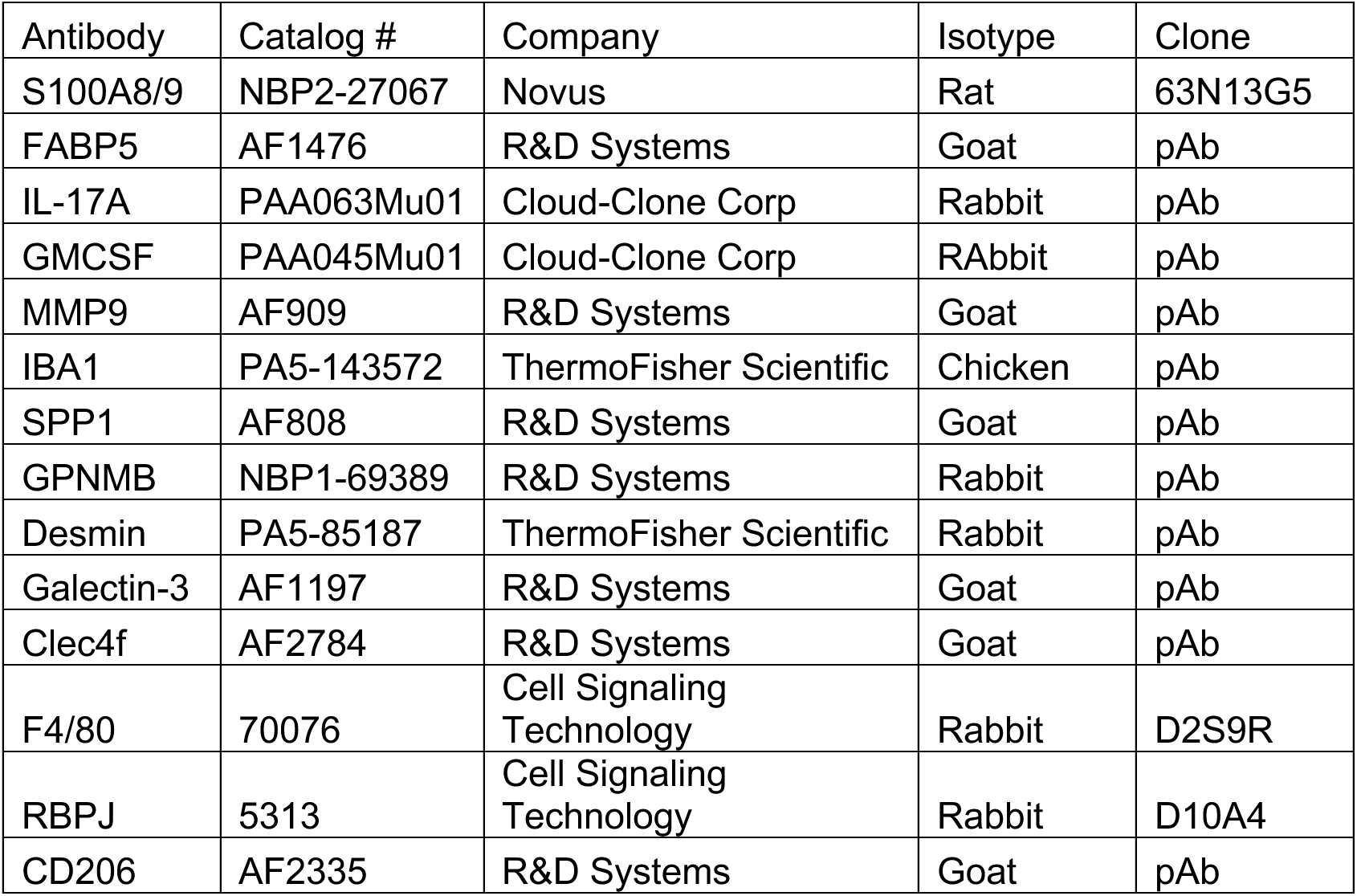
Mouse cycIF antibodies

## Notes

### Summary of Updates

Samir Lal who performed all bulk-RNAseq analysis in Figure 1 and Figure 2 was missing from the Author list. His work is a significant contribution to this manuscript.

